# Minimal *in vivo* requirements for developmentally regulated cardiac long intergenic non-coding RNAs

**DOI:** 10.1101/793257

**Authors:** Matthew R. George, Qiming Duan, Abigail Nagle, Irfan S. Kathiriya, Yu Huang, Kavitha Rao, Saptarsi M. Haldar, Benoit G. Bruneau

## Abstract

Long intergenic non-coding RNAs (lincRNAs) have been implicated in aspects of gene regulation, but their requirement for development needs empirical interrogation. To begin to understand the roles lincRNAs might play in heart development, we computationally identified nine murine lincRNAs that have developmentally regulated transcriptional and epigenomic profiles specific to early heart differentiation. Six of the nine lincRNAs had in vivo expression patterns supporting a potential function in heart development, including a transcript downstream of the cardiac transcription factor *Hand2* that we named *Handlr (<u>Hand</u>2*-associated linc<u>R</u>NA), *Rubie,* and *Atcayos*. We genetically ablated these six lincRNAs in mouse, which implicated genomic regulatory roles to four of the cohort, However, none of the lincRNA deletions led to severe cardiac phenotypes. Thus, we stressed the hearts of adult *Handlr* and *Atcayos* mutant mice by transverse aortic banding and found that absence of these lincRNAs did not affect cardiac hypertrophy or left ventricular function post-stress. Our results support roles for lincRNA transcripts and/or transcription to regulation of topologically associated genes. However, the individual importance of developmentally-specific lincRNAs is yet to be established. Their status as either gene-like entities or epigenetic components of the nucleus should be further considered.

## Introduction

A substantial portion of the mammalian genome is transcribed throughout development, while only a small fraction of this yields functional protein (Wong, Passey et al. 2001, 2012, Hon, Ramilowski et al. 2017). The remaining noncoding RNA is arbitrarily classified into long (lncRNA) and short transcripts based upon length greater or less than 200nt. A few lncRNAs have been implicated to be important for cardiac development (Grote, Wittler et al. 2013, Han, Li et al. 2014, Kurian, Aguirre et al. 2015, Anderson, Anderson et al. 2016). However, these RNA molecules are often products of the pervasive bidirectional transcription taking place at most genes (Katayama, Tomaru et al. 2005), which makes independently dissecting their function difficult. On the other hand, thousands of putative intergenic lncRNAs (lincRNAs) with little protein coding potential exist as stand-alone units (Carninci, Kasukawa et al. 2005). They can exhibit characteristics indicative of epigenetic control, such as histone H3 trimethylation at lysine 4 (H3K4me3) and acetylation of lysine 27 (H3K27Ac) at promoters and trimethylation at lysine 36 (H3K36me3) throughout their gene body, splicing, 5’ m7G capping, and polyadenylation (Derrien, Johnson et al. 2012, Sati, Ghosh et al. 2012, Quinn and Chang 2016). LincRNAs also can display considerable sequence conservation and are dynamically expressed in specific tissues at developmentally discrete times (Diederichs 2014, Perry and Ulitsky 2016, Mattioli, Volders et al. 2019). For example, the lincRNAs *Braveheart*, *Meteor, and Carmen* seem to play significant roles in precardiac mesodermal differentiation, at least in cellular differentiation systems *in vitro* (Klattenhoff, Scheuermann et al. 2013, Ounzain, Micheletti et al. 2015, Alexanian, Maric et al. 2017, Hou, Long et al. 2017). However, *Meteor* knockout *in vivo* resulted in milder phenotypes than observed *in vitro* (Guo, Xu et al. 2018), while *Braveheart* and *Carmen* have not been tested in embryos. Thus, few lincRNAs have been shown to be required for cardiac development *in vivo*. The energy investment a cell puts toward the processing and maintenance of these transcripts suggests their putative importance to the cell and organism. Therefore, efforts must be taken to interrogate specific lincRNA requirements for proper embryogenesis.

We were most interested in lincRNAs that might act to influence the early commitment of nascent mesoderm into the cardiac lineage. We hypothesized that as-yet unstudied transcripts were important for this most fundamental stage of cardiac development. Therefore, we screened for the expression of candidates during mouse embryonic stem cell (mESC) *in vitro* differentiation into cardiomyocytes (CM) through nascent mesoderm (MES), cardiac mesoderm (cMES), and cardiac progenitor (CP) intermediates. Of more than 114,000 long noncoding RNA annotations, we identified a small cohort of lincRNAs with epigenetic regulation, clear splice structure, and cardiac progenitor specificity, which we then validated *in vivo* in the early embryo. Ablation of these noncoding genes revealed regulatory roles within their topologically associated domains (TADs) but very mild or undetectable phenotypes in heart development or postnatal function.

## Results

### LincRNAs with cardiac-specific expression and epigenetic regulation in vitro

We hypothesized that, like many canonical genes, a subset of lincRNAs would be specifically expressed in the cardiac lineage. We also predicted that those most critical for heart formation would function early in its development. To find candidate lincRNAs, we re-mapped stranded raw RNA-seq reads from differentiations of mouse ESCs into cardiomyocytes (Figure 1A/B; (Wamstad, Alexander et al. 2012, Devine, Wythe et al. 2014) against Noncode version 4.0-annotated transcripts (Xie, Yuan et al. 2014). Additionally, we integrated parallel histone modification ChIP-seq data (Wamstad, Alexander et al. 2012) to discriminate loci under epigenetic regulation. To screen for cardiac developmental specificity, we chose to focus on elements that were lowly expressed in ESCs (FPKM < 0.5), while strongly upregulated in cardiac mesoderm or cardiac progenitors (FPKM >1.0). The majority of protein coding genes display antisense transcription from their promoters (Carninci, Kasukawa et al. 2005), including numerous studied lncRNAs (Kino, Hurt et al. 2010, Grote, Wittler et al. 2013, Han, Li et al. 2014, Kurian, Aguirre et al. 2015, Ramos, Andersen et al. 2015, Anderson, Anderson et al. 2016, Daneshvar, Pondick et al. 2016, Gore-Panter, Hsu et al. 2016, Xu, Zhang et al. 2016). We focused instead on genomic elements that could be altered independently from their nearby protein-coding genes. Therefore, we filtered for RNA annotations whose transcriptional start site (TSS) began more than 1 kilobase (kb) from the TSS of known protein-coding genes. To avoid spurious transcripts, we required candidates be spliced and then further refined the list to those displaying histone H3 lysine-4 trimethylation (H3K4me3) and H3 lysine-27 acetylation (H3K27Ac) at their promoters. After removing annotated transcripts that splice into nearby protein coding genes (i.e. A930006K02Rik into *Ifnar1*), we found that these criteria narrowed candidates to only nine total lincRNAs out of 114,104 considered transcripts (Figure 1B).

**Figure 1.**
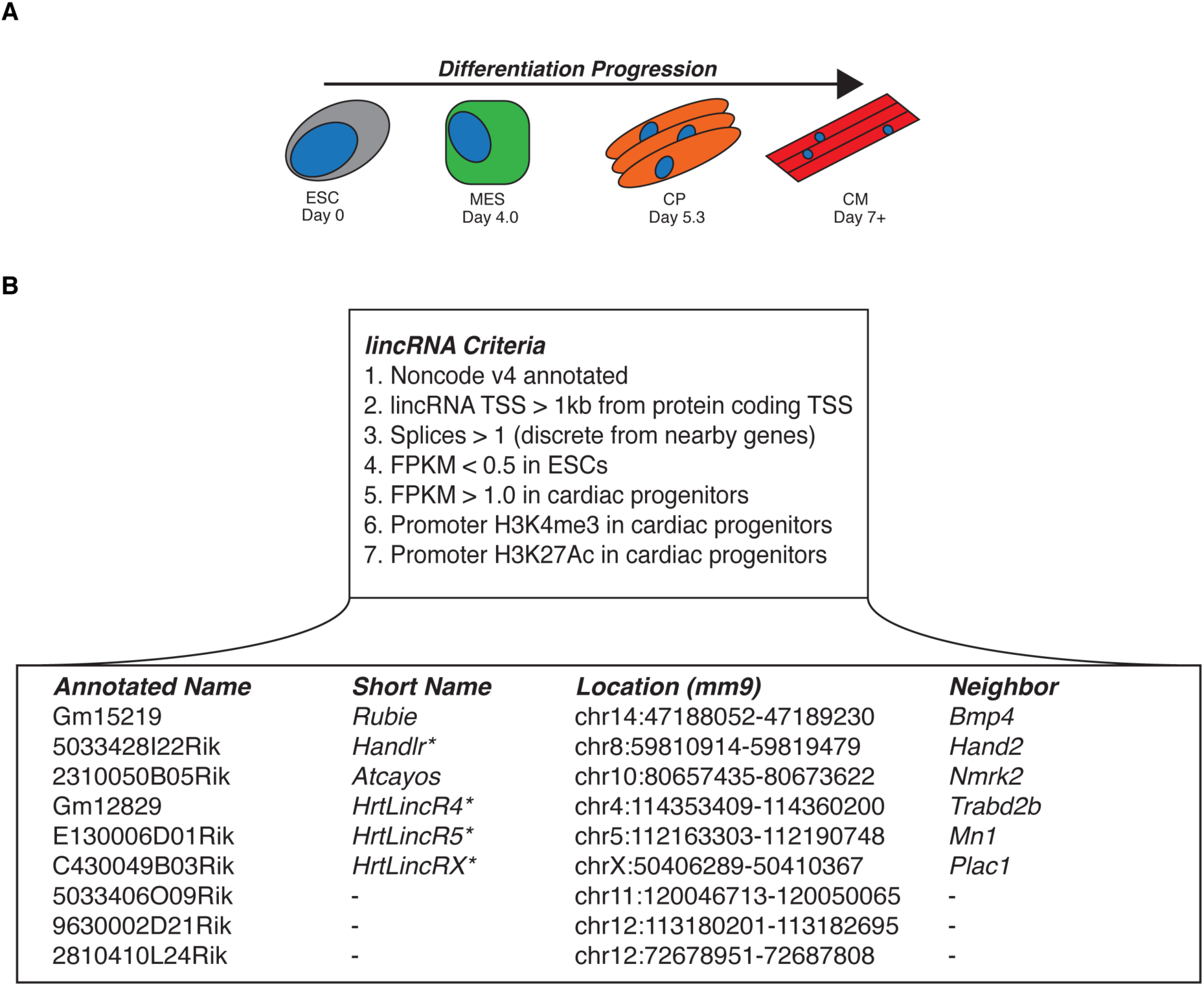
Epigenetically regulated cardiac lincRNAs. A. Differentiation progression of mESCs into cardiomyocytes used for lincRNA candidate selection; ESC, embryonic stem cell, MES, mesoderm, CP, cardiac progenitor, CM, cardiomyocyte. B. Criteria for lincRNA identification and resulting 9 candidates; *, name assigned by Bruneau lab.

The lincRNA *Rubie* (**R**na **U**pstream **B**mp4 in the **I**nner **E**ar, Gm15219) was known to co-express with *Bmp4* after E15.0 in the mouse inner ear, and its perturbed splicing was previously implicated in ear vestibule malformation and consequent circling behavior (Roberts, Abraira et al. 2012). However, our candidate screen revealed it to be expressed much earlier in the developing cardiac mesoderm(Figure 2A). As in the inner ear, it’s expression *in vitro* overlapped the TGF-β signaling protein *Bmp4* (Wozney, Rosen et al. 1988) and these genes, separated by approximately 176kb, co-occupied a strongly interacting region within the same topologically associated domain (TAD; Figure 2C) (Dixon, Selvaraj et al. 2012, Nora, Lajoie et al. 2012).

**Figure 2.**
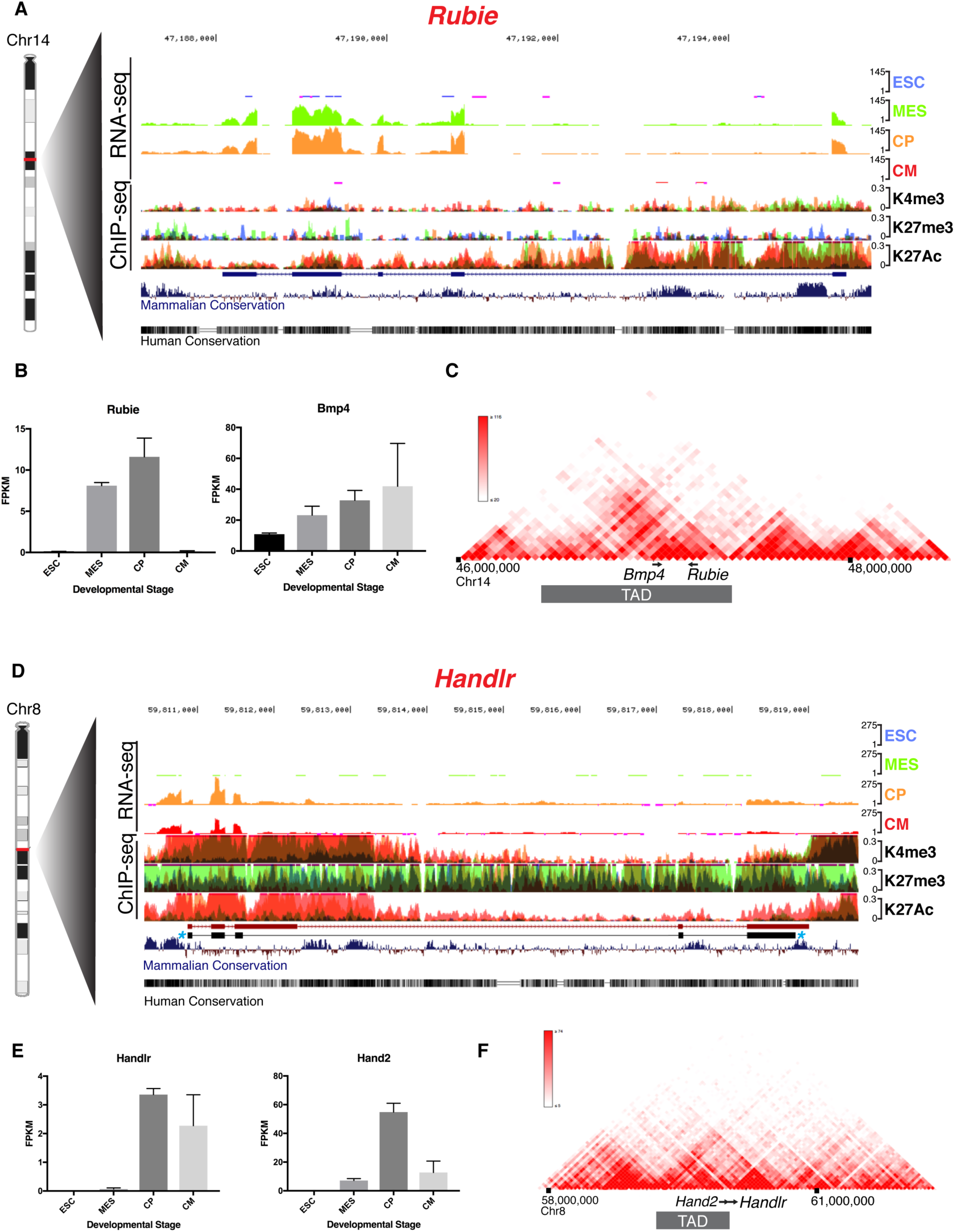
Genomic characterization of *Rubie* and *Handlr in vitro.* A. UCSC Genome Browser tracks of *Rubie* RNA-seq and overlaid histone H3 ChIP-seq at ESC, MES, CP, and CM stages of *in vitro* differentiation; RefSeq annotation in blue. B. Quantified expression of *Rubie* and *Bmp4* at each differentiation stage. C. 3D Genome Browser Hi-C heatmap of chromosome interactions around *Bmp4* and *Rubie* loci. D. UCSC Genome Browser tracks of *Handlr* RNA-seq and overlaid histone H3 ChIP-seq at ESC, MES, CP, and CM stages of *in vitro differentiation*. Ensembl annotation in red; actual exon structure of predominant *Handlr* transcript in black with blue stars. E. Quantified expression of Handlr and Hand2 at each differentiation stage. F. 3D Genome Browser Hi-C heatmap of chromosome interactions around *Handlr* and *Hand2* loci; TAD, topologically associated domain. ESC, embryonic stem cell, MES, mesoderm, CP, cardiac progenitor, CM, cardiomyocyte; blue, ESC, green, MES, orange, CP, red, CM; K4me3, histone H3 lysine 4 trimethylation; K27me3, histone H3 lysine 27 trimethylation; K27Ac, histone H3 lysine 27 acetylation; TAD, topologically associated domain.

*Hand2*, a transcription factor critical for heart development (Srivastava, Thomas et al. 1997), was previously shown to be regulated by antisense transcription of the noncoding RNA *Upperhand* (*Uph*) locus 5’ from its promoter (Anderson, Anderson et al. 2016). Our search identified *5033428l22Rik* as a candidate lincRNA approximately 8 kb downstream of *Hand2*, which we named *Handlr* (<u>Hand</u>2-Associated linc<u>R</u>NA). During the course of our study, others also studied this gene in the cardiac lineage, which they named *Handsdown* (*Hdn*) (Ritter, Ali et al. 2019). This region displayed numerous transcribed splice forms, but 3’ rapid amplification of cDNA ends (RACE) of E9.5 cDNA revealed a single predominant 5-exon, polyadenylated isoform that varied from its Ensembl- and RefSeq-annotated structures (Figure 1D, Supplemental figure S1A, S1B). *Handlr’s* expression overlapped *Hand2 in vitro* (Figure 2E), but these genes sat near a TAD border (Figure 2F) and were separated by a CTCF insulation site (Martin, Pantoja et al. 2011), suggesting a potential topological division between the two.

Seven additional annotated lincRNAs met the criteria for subsequent analyses. We discovered *Atcayos* (*2310050B05Rik*) transcription to span the important cardiomyocyte metabolic regulator *Nmrk2* (Diguet, Trammell et al. 2018) and precede its expression in differentiating cardiac progenitors and cardiomyocytes (Supplemental figure S2A, S2B). *Gm12829*, named *HrtLincR4* (<u>h</u>ea<u>rt</u> <u>lincR</u>NA of chromosome <u>4</u>) was correlatively expressed within a genomic domain in frequent contact with *Trabd2b*, a Wnt protein-binding metalloprotease (Zhang, Abreu et al. 2012). In addition, its expression was only transiently detected within an 18-hour window at the cardiac mesoderm (cMES) stage of differentiation (Supplemental figure S2C-S2E). Also, *E130006D01Rik*, named *HrtLincR5* (<u>h</u>ea<u>rt</u> <u>lincR</u>NA of chromosome <u>5</u>), was expressed within a *Mn1*-interacting DNA domain approximately 275kb downstream of this transcriptional coactivator (van Wely, Molijn et al. 2003). This transcript displayed highly stereotypic splicing and was only detected at the cardiac progenitor stage of differentiation (Supplemental figure S3A-S3C). *C430049B03Rik*, named *HrtLincRX* (<u>h</u>ea<u>rt</u> <u>lincR</u>NA of <u>X</u> chromosome) was highly expressed early in our differentiation model and contained a miRNA cluster in its 3’ tail that had previously been shown to drive cardiomyocyte specification (Shen, Soibam et al. 2016). This lincRNA also lies approximately 12.5kb downstream of- and overlapped expression with-the essential placental gene *Plac1* (Jackman, Kong et al. 2012) not normally expressed in most somatic tissues (Fant, Farina et al. 2010) (Supplemental figure S2D-S2F). Finally, *5033406O09Rik*, *9630002D21Rik*, and *2810410L24Rik* also fulfilled the criteria of our screen (Supplemental figure S4A-S4C).

All nine lincRNAs contained regions with highly homologous sequence to human and/or mammalian genomes (Figure 2, Supplemental figures S2-S4). To assess the protein coding potential of these candidates, we employed multiple tests. First, we evaluated PhyloCSF (Lin, Jungreis et al. 2011) codon scores in all three frames for each transcript. Whereas this algorithm readily detected stretches with coding potential in known genes such as *Bmp4* and the micropeptide-containing *Apela/Toddler* (Pauli, Norris et al. 2014) and *Dworf* (Nelson, Makarewich et al. 2016), we found no evidence for protein coding potential in our lincRNA cohort, with one exception. A 28 amino acid reading frame in the second exon of *HrtLincR4* was predicted to represent a possible conserved coding region (Supplemental figure S5A), though HrtLincR4, as well as each additional member of the cohort displayed negative coding-non-coding indices (CNCI, Supplemental figure S5B) (Sun, Luo et al. 2013) similar to the known paraspeckle-associated lincRNA Neat1 (Hutchinson, Ensminger et al. 2007). However, CPAT (Wang, Park et al. 2013) calculation of hexamer usage bias (Wang, Park et al. 2013) and Ficket nucleotide composition and codon usage bias (Fickett 1982) could not differentiate our lincRNA group from *Apela* or *Dworf* (Supplemental figure S5C). We next tested *Rubie*, *Handlr*, *Atcayos*, *HrtLincR4*, *HrtLincR5*, and *HrtLincRX* localization in fractionated cardiac progenitor cells. These lincRNAs were biased to the nucleus, where *Rubie* and *Handlr* were enriched even more so than *Neat1* (Supplemental figure S5D). Additionally, these six lincRNAs could generate cDNA using oligo dT primers at least as efficiently as *Actb* and *Neat1*, known to be polyadenylated (Sasaki, Ideue et al. 2009) (Supplemental figure S5E, Supplemental table S1). Taken together, we classified these lincRNAs as nuclear-enriched, polyadenylated, transcripts with little translational capacity. However, we could not rule out the coding potential of the minority *HrtLincR4* fraction that reaches the cytoplasm.

### A cohort of screened cardiac lincRNAs display dynamic expression in vivo in the developing mouse heart

We examined the spatiotemporal expression patterns in the developing embryo for each of the nine candidate lincRNAs by whole mount *in situ* hybridization from E7.25 through E10.5 using transcript-specific probes (Supplemental table S2). The expression patterns observed *in vitro* were largely predictive of those observed *in vivo. Rubie* was first observed in the E7.75 embryo, where, similar to *Bmp4 (Perea-Gomez, Shawlot et al. 1999),* it strongly demarcated the extraembryonic boundary and flanked the eventual heart field. From E8.0 to E8.5, its expression became less focused, spreading throughout the developing cardiac crescent and heart tube, respectively. *Rubie* transcription at E8.75 was largely non-cardiac, and by E9.5, it was strongly localized to posterior mesoderm and the otic vesicle (Figure 3A). This patterning overlapped a somewhat refined subset of what had previously been established for Bmp4 (Danesh, Villasenor et al. 2009).

**Figure 3.**
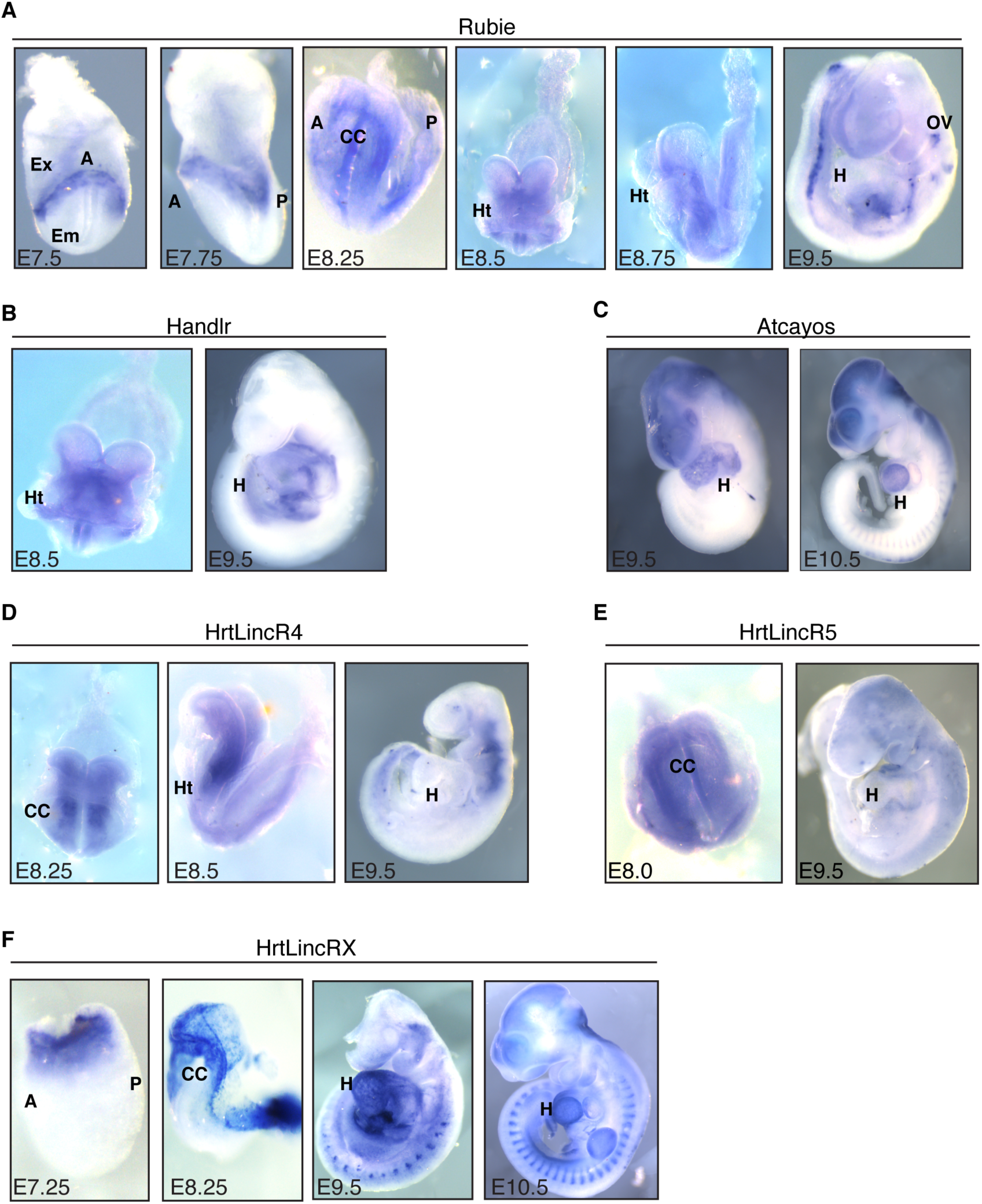
lincRNA expression patterns *in vivo*. A. *in situ* hybridization staining for *Rubie* from E7.5 through E9.5. B. *in situ* hybridization staining for *Handlr* at E8.5 and E9.5. C. *in situ* hybridization staining for *Atcayos* at E9.5 and E10.5. D. *in situ* hybridization staining for *HrtLincR4* at E8.25, E8.5, and E9.5. E. *in situ* hybridization staining for *HrtLincR5* at E8.0 and E9.5. F. *in situ* hybridization staining for *HrtLincRX* from E7.5 through E10.5. A, anterior; P, posterior; Em, embryonic region; Ex, extraembryonic region; CC, cardiac crescent; Ht, heart tube; H, heart; OV, otic vesicle.

From E8.5 through E9.5, *Handlr* was transcribed in the developing heart tube, posterior cardiac progenitors, branchial arches, and lateral plate mesoderm (Figure 3B). These patterns overlapped what was also shown for *Hand2* at this developmental stage (Charite, McFadden et al. 2000), suggesting common regulation between *Hand2* and *Handlr*. *Atcayos*, as predicted by *in vitro* expression patterns, was weakly expressed during early stages of heart tube formation, while it was dramatically upregulated after E9.5 in the developing ventricles, as well as cranial structures and somitic mesenchyme (Figure 3C).

From E8.25 through E9.5, *HrtLincR4* displayed strong expression in developing pharyngeal mesoderm, just dorsal to the developing cardiac crescent (Figure 3D). Given its highly transient expression within differentiating cardiac mesoderm *in vitro*, these data suggested *HrtLincR4* to be quickly specified to the secondary heart field and/or adjacent tissues during the onset of cardiac lineage commitment. *HrtLincR5* was broadly expressed throughout the mesoderm, including the nascent cardiac crescent, at E8.25. As expected by its short-lived *in vitro* expression pattern, *HrtLincR5* was predominantly lost *in vivo* by E9.5 (Figure 3E).

*HrtLincRX* was strongly expressed by E7.5 during cardiac lineage formation in anterior mesoderm at the extraembryonic boarder, as well as in extraembryonic tissues. At E8.25, it was strongly expressed in the cardiac crescent, amniotic membranes, and the developing allantois. While expression of the adjacent *miR322*/*503* cluster was previously shown to be cardiac-specific (Shen, Soibam et al. 2016), this lincRNA was widely expressed throughout the heart, forelimb, and somitic mesoderm at E9.5 and E10.5 (Figure 3F). This suggested divergent regulation and/or compounding roles for *HrtLincRX* versus its miRNA components. We could not effectively validate the expression of *5033406O09Rik*, *9630002D21Rik*, or *2810410L24Rik* beyond diffuse, low levels in the developing mouse embryo (Supplemental figure S6A-S6C). These experiments established the striking expression patterns of numerous tissue-specific lincRNAs identified from our screen of *in vitro* cardiac differentiation. Therefore, we aimed to test developmental importance of *Rubie, Handlr, Atcayos, HrtLincR4, HrtLincR5, and HrtLincRX* expression during embryonic development.

### Cas9 ablation of lincRNA promoter regions in vivo identifies local gene regulatory roles

To determine the requirement for the six lincRNAs that displayed compelling *in vivo* expression, we generated knockout mouse lines through pronuclear Cas9 mRNA and tru-sgRNA (Fu, Sander et al. 2014) injections. For each knockout, paired tru-sgRNAs were co-injected to induce 2-3kb deletions flanking the respective lincRNA transcriptional start site (TSS) (Figure 4A, Supplemental table S3), which successfully generated heritable alleles for all six target regions. After substantial outbreeding (> 3 backcrosses into C57Bl/6j background), we crossed heterozygotes and harvested the anterior half of E8.25 embryos for RT-qPCR (Figure 4B). We found that these deletions ablated downstream transcription of each lincRNA (Figure 4C-4H). As these lincRNAs were nuclear-enriched, we hypothesized they might be involved in transcriptional regulation within their local genomic environments. To test this, we measured expression of neighboring protein-coding genes sharing the same respective TADs (Supplemental table S1). While *Rubie* was previously associated with BMP4 signaling in the inner ear (Roberts, Abraira et al. 2012), its requirement for *Bmp4* expression had not been established. We found that loss of *Rubie* resulted in significant reduction of *Bmp4* expression during cardiac specification. Furthermore, the amount of transcribed *Rubie* was directly correlated with *Bmp4* levels in this region at the same time point. This effect was maintained even within equivalent underlying genotypes, whereby *Rubie* and *Bmp4* transcript levels were still significantly correlated among *Rubie*^+/-^ offspring only (Figure 4C). These data strongly suggested that either the act of *Rubie* transcription and/or its physical RNA molecule were responsible for quantitative regulation of *Bmp4* expression.

**Figure 4.**
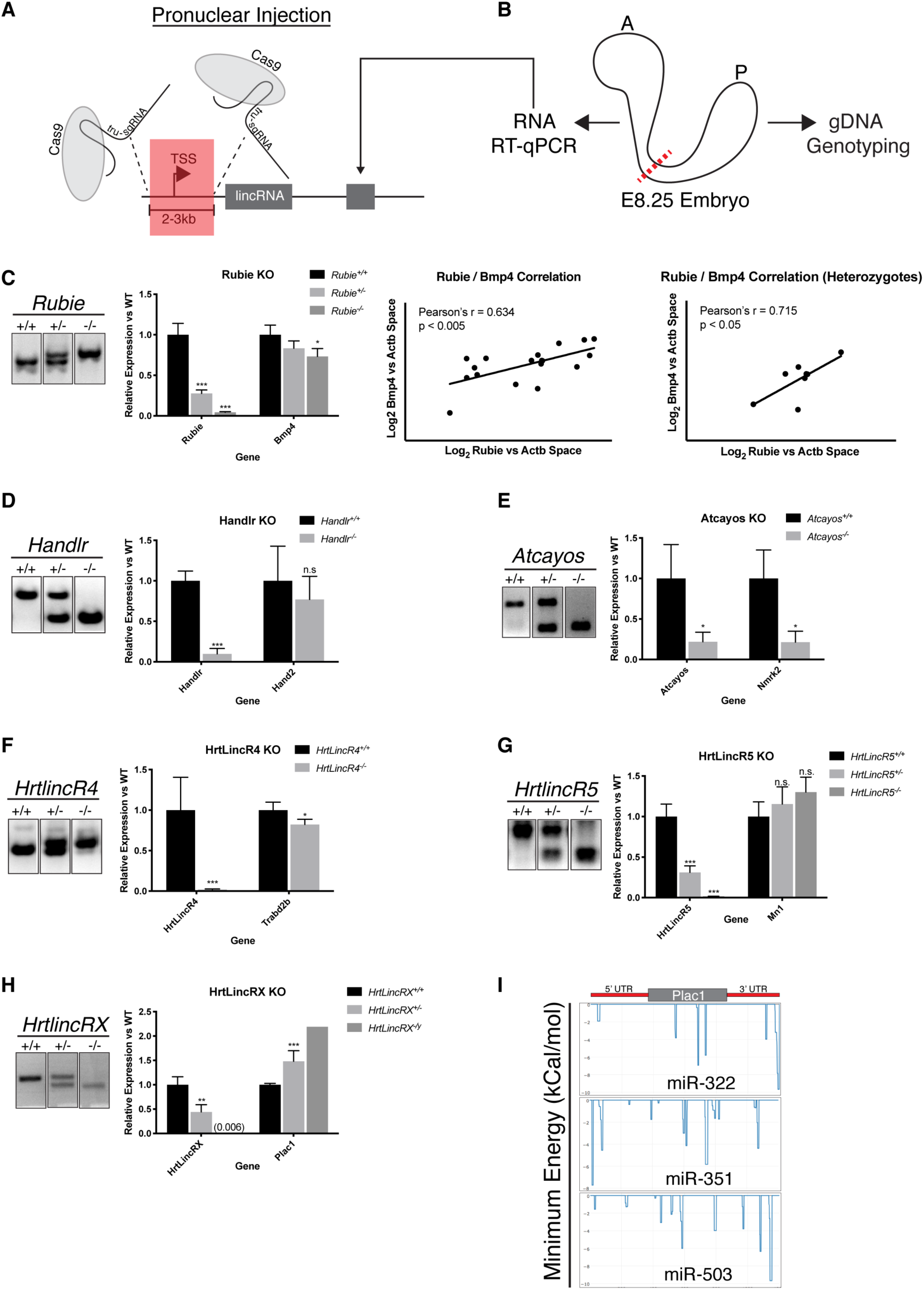
Cas9 ablation of cardiac lincRNAs *in vivo* and effects on local gene expression. A. lincRNA TSS/promoter ablation strategy; TSS, transcriptional start site; tru-sgRNA, truncated single guide RNA. B. Schematic for RT-qPCR on anterior half of E8.25 embryo; A, anterior, P, posterior, red line, bisection point. C. Left: Electrophoresed gDNA PCR genotyping products of *Rubie* alleles and resulting *Rubie* and *Bmp4* expression in anterior E8.25 embryos; Right: correlation between *Rubie* expression and *Bmp4* expression for all genotypes or *Rubie*^+/-^ only, respectively. D. Electrophoresed gDNA PCR genotyping products of *Handlr* alleles and resulting *Handlr* and *Hand2* expression in anterior E8.25 embryos. E. Electrophoresed gDNA PCR genotyping products of *Atcayos* alleles and resulting *Atcayos* and *Nmrk2* expression in anterior E8.25 embryo. F. Electrophoresed gDNA PCR genotyping products of *HrtLincR5* alleles and resulting *HrtLincR5* and *Mn1* expression in anterior E8.25 embryos. G. Electrophoresed gDNA PCR genotyping products of *HrtLincR4* alleles and resulting *HrtLincR4* and *Trabd2b* expression in anterior E8.25 embryos. H. Left: Electrophoresed gDNA PCR genotyping products of *HrtLincRX* alleles and resulting *HrtLincRX* and *Plac1* expression in anterior E8.25 embryos; Right: IntaRNA 2.0 binding prediction between *HrtLincRX* 3’ miRNAs and Plac1. *, p < 0.05; ***, p < 0.005; n.s., not significant; Student’s 2-tailed t-test. Data presented as mean +/-SEM.

Despite proximity to and co-expression with *Handlr*, *Hand2* activation was not dependent on *Handlr* lincRNA (or its underlying promoter DNA sequence, Figure 4D). We speculated that the TAD architecture and CTCF boundary between these genes may introduce complex dynamics within the region. We also could not find a correlation between *Mn1*’s expression to *HrtLincR5* (Figure 4E). In contrast, *Nmrk2* and *Trabd2b* expression was dependent on *Atcayos* (Figure 4F) and *HrtLincR4* (Figure 4G), respectively. Furthermore, *Plac1* transcription was significantly and inversely correlated to *HrtLincRX* levels, whereby loss of *HrtLincRX* resulted in approximately a 2-fold increased expression of *Plac1* (Figure 4H). However, using IntaRNA (Mann, Wright et al. 2017) analysis, we calculated stable RNA-RNA interactions between all three miRNAs constituents of its 3’ tail (miRNA-322, miRNA-351, miRNA-503) and the 5’- and 3’-untranslated regions (UTRs) of *Plac1* (Figure 4I). Therefore, this relationship could likely be explained by the loss of inhibitory miRNA binding to *Plac1* primary transcript. Nonetheless, these data indicated a potential role for *HrtLincRX* and/or its miRNAs in demarcating embryonic from extraembryonic mesoderm during gastrulation and early cardiogenesis.

### Cardiac lincRNAs are not required for viable mouse development

To determine the requirement of our lincRNA cohort for viable embryonic development *in vivo*, we bred heterozygotes for each gene and examined ratios of expected offspring that survived to weaning (Supplemental table S4). We could not establish any reduction in viability within null progeny for any tested lincRNA (Figure 5A, 5C, 5D, Supplemental figure S7A-S7C). Furthermore, homozygous offspring for these loci lived to adulthood and were fertile. However, the *Rubie* null genotype did sporadically recapitulate the circling behavior described by Roberts et al, which they observed as a result of aberrant *Rubie* splicing in the SWR/J genetic background (Roberts, Abraira et al. 2012). While rarely observed, circling was only present in *Rubie*^-/-^ mice during >2 years of colony breeding (Figure 5B). Nonetheless, despite their clear expression within the developing heart, we concluded that none of the lincRNAs were individually required for viable development or fertility in the FVBn; C57BL/6j mixed background.

**Figure 5.**
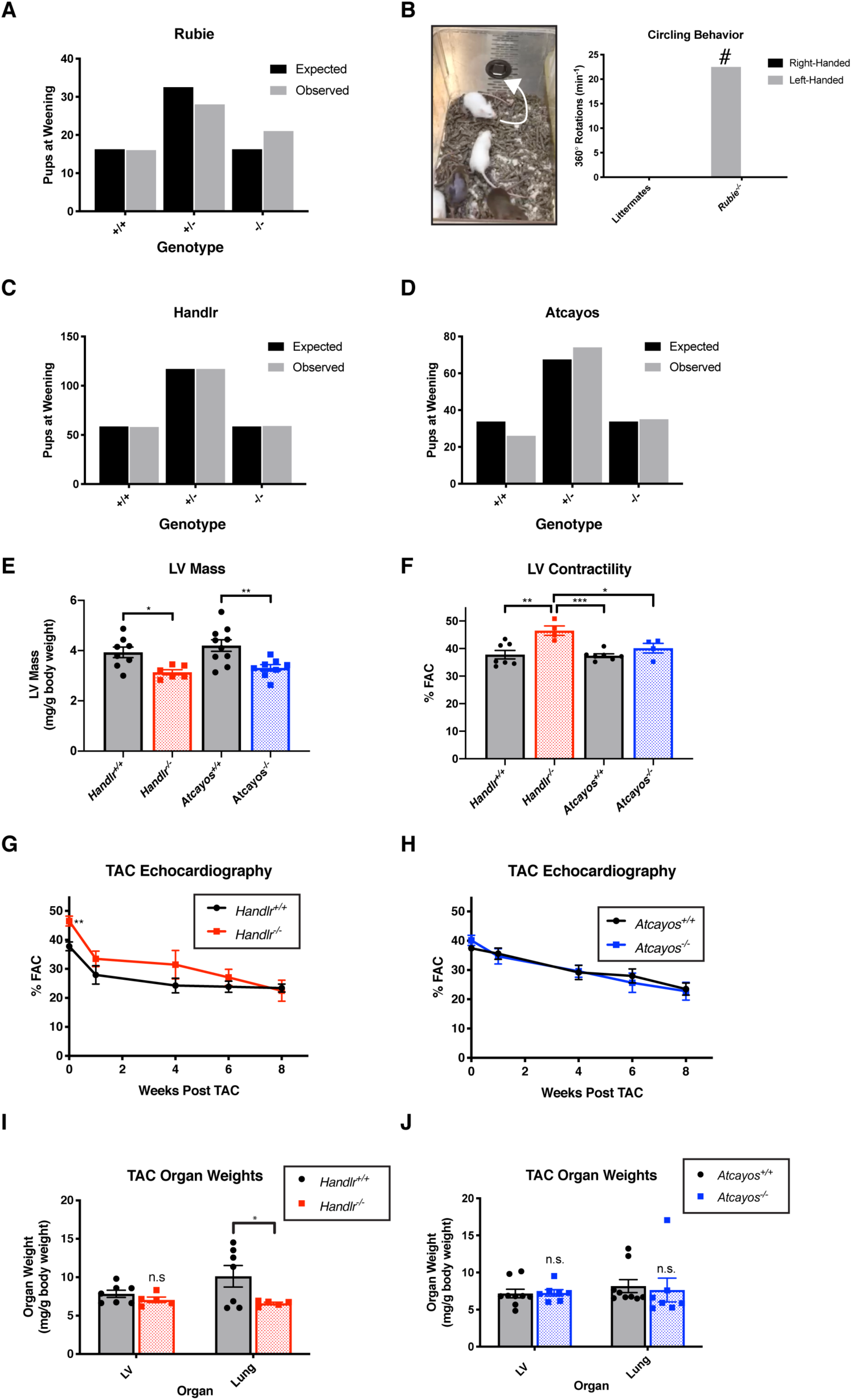
Viability and phenotypic effects after lincRNA knockout. A. Offspring recovered at weaning from *Rubie^+/-^* x *Rubie^+/-^*cross vs expected Mendelian ratios. B. Representative sporadic circling behavior in *Rubie^-/-^* offspring; #, only observed in *Rubie^-/-^*genotype over 2+ years of observation. C. Offspring recovered at weaning from *Handlr^+/-^* x *Handlr^+/-^* cross vs expected Mendelian ratios. D. Offspring recovered at weaning from *Atcayos^+/-^*x *Atcayos^+/-^* cross vs expected Mendelian ratios. E. LV mass calculated by echocardiographic measurements in *Handlr* WT versus NULL and *Atcayos* WT versus NULL litter-matched adult males, respectively. F. LV contractility measured by echocardiography in *Handlr* WT versus NULL and *Atcayos* WT versus NULL litter-matched adult males, respectively. G. Time course of cardiac contractility after TAC in *Handlr* WT versus NULL litter-matched adult males. H. Time course of cardiac contractility after TAC in *Atcayos* WT versus NULL litter-matched adult males. I. Week 8 post-TAC LV and lung weights in *Handlr* WT versus NULL litter-matched adult males. J. Week 8 post-TAC LV and lung weights in *Atcayos* WT versus NULL litter-matched adult males. *, p <0.05; **, p < 0.01; ***, p <0.005, Student’s 2-tailed t-test, except panel I, Z-test; LV, left ventricle; FAC, fractional area shortening; TAC, transverse aortic constriction; WT, wild type.

### Neither Handlr nor Atcayos play important roles in the cardiac stress response

Gene knockout models often require external stressors to materialize overt phenotypes. *Handlr* and *Atcayos* were the only cohort lincRNAs expressed in adult hearts, and *Atcayos’* very high expression was reduced approximately 50% after transverse aortic constriction (TAC)-induced cardiac hypertrophy (Supplemental figure S7D, S7E) (Duan, McMahon et al. 2017). Therefore, we performed TAC experiments on *Handlr-* and *Atcayos*-null mice and compared their responses to wild type (WT) littermates. At baseline, calculated left ventricular (LV) masses (from echocardiographic measurements) were modestly reduced in *Handlr^-/-^* and *Atcayos^-/-^* adults (-18.4%, p < 0.05; -22.4%, p < 0.01, respectively; Figure 5E; Supplemental table S5 and S6).

Additionally, *Handlr*-null adults had slightly but significantly increased fractional area contractility (FAC) over *Handlr^+/+^*, *Atcayos^+/+^,* and *Atcayos^-/-^* genotypes (46.5%, versus 37.8%, p < 0.01; 37.4%, p < 0.005; and 40.1%, p < 0.05, respectively; Figure 5F, Supplemental table S5/S6). However, neither loss of *Handlr* nor *Atcayos* could invoke a significant alteration to LV mass increase or LV fractional shortening decrease after TAC-mediated stress. Of note, the severity of the TAC-response in *Handlr*-related experiments was greater than that for *Atcayos* due to genetic background and/or surgical procedure differences for each cohort. This resulted in sharper fractional shortening decrease and longer duration under cardiac failure (%FAC < 30%) for these mice. Consequently, *Handlr^+/+^*males displayed greater downstream increase in lung mass versus *Handlr^-/-^*individuals (whom began the experiment with greater contractile function), while only sporadic lung hypertrophy arose in the *Atcayos* groups (Figure 5G-5J). This result was not interpreted to indicate an altered cardiac hypertrophic response in *Handlr-*null individuals but further supports their increased LV FAC measured at baseline. We also could not detect any noticeable changes versus WT in the expression of canonical hypertrophic response genes *Nppa*, *Nppb*, or *Acta1* due to *Handlr* nor *Atcayos* knockout (data not shown). Therefore, despite strong *Atcayos* expression in the adult heart and the known hypertrophic involvement of *Handlr*’s neighbor *Hand2*, loss of these transcripts did not induce an altered response to LV pressure overload. Therefore, we concluded that neither of these lincRNAs played important roles in the physiological response to heart failure.

### Compound heterozygosity reveals genetic interaction between Rubie and Bmp4 but not between Handlr and Hand2

Despite the lack of overt lethality in lincRNA-deficient offspring, we carefully examined morphological heart development in *Handlr* and *Rubie* null embryos, the only conditions that produced noticeable physiological effects. We harvested E15.5 hearts and examined transverse histological sections to establish any change to chamber septation, myocardial trabeculation and/or compaction, or ventricular outflow tract (OFT) development. *Handlr*^-/-^ adults exhibited increased left ventricular fractional shortening over WT controls, but we could not associate this functionality with overt changes in cardiac anatomy (Supplemental figure S8A, S8B). Due to overlapping expression patterns between *Hand2* and *Handlr* in the developing heart, we next tested embryos from *Hand2^+/-^ x Handlr^+/-^* crosses to eliminate one allele of either *Hand2* or *Handlr* per chromosome. However, neither *Hand2* heterozygosity nor *Hand2^+/-^; Handlr ^+/-^* compound heterozygosity resulted in any clear effects on heart morphogenesis (Supplemental figure S8C, S8D). In addition, we did not notice any elevated lethality in *Hand2^+/-^;Handlr ^+/-^* offspring (data not shown, n = 63).

*Bmp4* expression in the embryo was correlated to the amount of *Rubie* transcript. Numerous studies have established the requirement of proper BMP4 dosage for normal septation of the atria, ventricles, and outflow tract (OFT), as well as viable embryo development (Dunn, Winnier et al. 1997, Jiao, Kulessa et al. 2003, Goldman, Donley et al. 2009). Therefore, we tested the hypothesis that compound haploidy of *Bmp4* and *Rubie* together would result in an exacerbated onset of resulting phenotypes. For this, we bred *Bmp4^fl/fl^ x Rubie^+/-^*; *Actb-Cre^+^* (single transgene integration) in the FVB/n; C57BL/6j mixed genetic background. When we examined E15.5 hearts, neither the *Rubie^-/-^* background nor loss of a single *Bmp4* allele could induce an abnormal cardiac phenotype. We also recovered these offspring in expected ratios at weaning (Figure 6A-6D). However, we did find a sustained approximately 20% reduction in recovered pups carrying the *Bmp4^+/-^/Rubie^+/-^* compound genotype (Figure 6D; n = 186). In addition, these offspring exhibited incidences of OFT distortion out of the right ventricle beyond its typical boundary. In these cases, the origins of the pulmonary artery skewed toward the left ventricular OFT and aortic valve (Figure 6E). While we were unable to clearly establish communication between the pulmonary and aortic outflow systems in these instances, the data point toward a modest genetic interaction between *Rubie* and *Bmp4* in cardiac morphogenesis.

**Figure 6.**
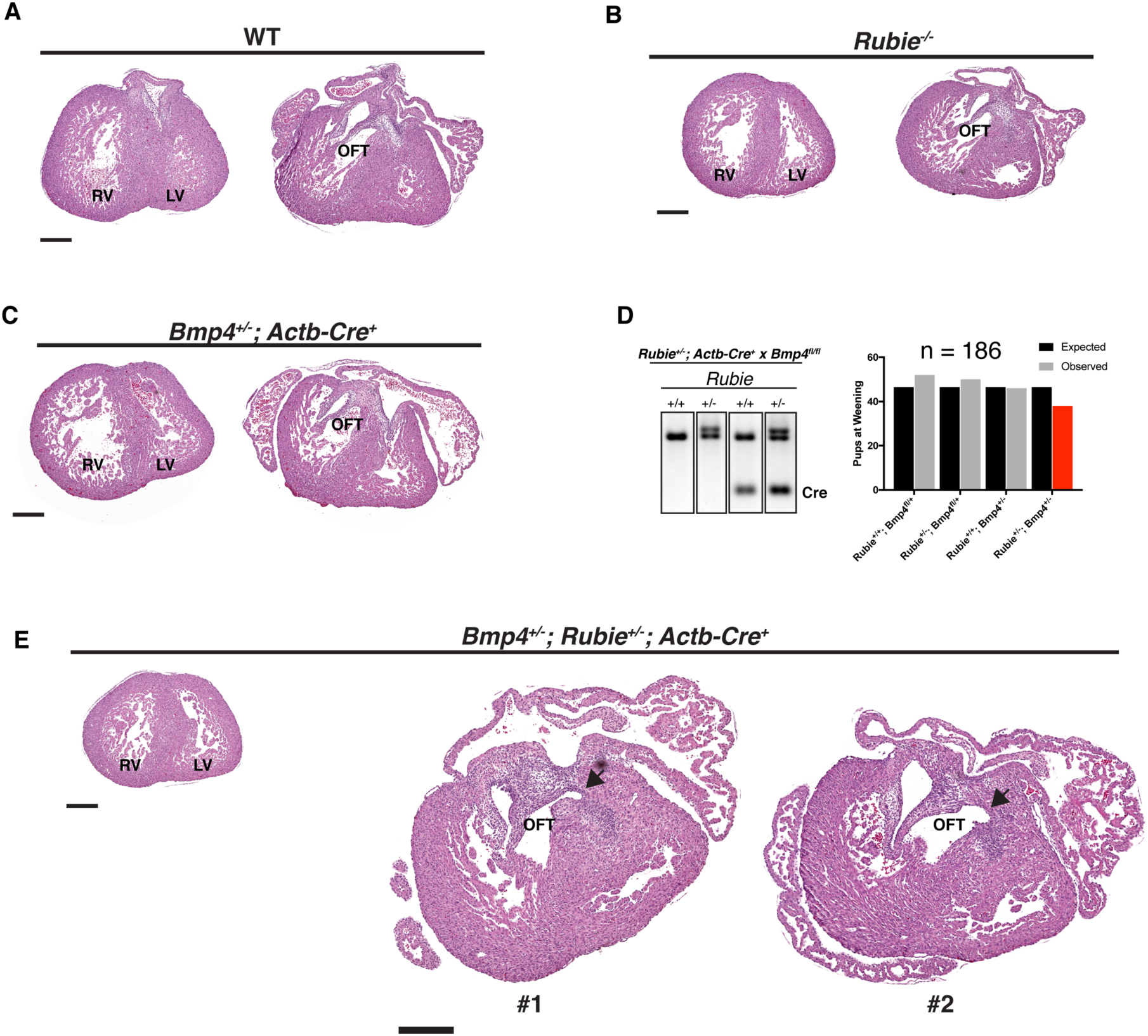
Effect of *Rubie* ablation on heart development at E15.5. A-C,E. Oblique transverse hematoxylin and eosin histological sections of cardiac ventricular and OFT morphogenesis, respectively, at E15.5. A. Representative wild type WT morphology. B. Representative Rubie^-/-^ morphology. C. Representative *Bmp4^+/-^* (*Actb-Cre^+^*) morphology. D. Left: Electrophoresed gDNA PCR genotyping products of *Rubie* and *Actb-Cre* transgene alleles, respectively. Right: Offspring recovered at weaning from *Rubie^+/-^*; *Actb-Cre^+^* x *Bmp4^fl/fl^*mating vs expected Mendelian ratios G. H. Representative *Bmp4^+/-^*; *Rubie^+/-^* (*Actb-Cre^+^*) morphology in 2 separate individuals. RV, right ventricle; LV, left ventricle; OFT, outflow tract; WT, wild type; scale bar, 300μm; arrow, distorted OFT orientation.

## Discussion

With thousands of uncharacterized noncoding transcriptional elements expressed throughout the genome, efforts must be taken to better understand the functional relevance of unstudied lincRNAs. Towards this need, these experiments were meant to identify and test the requirement for cardiac progenitor-specific lincRNAs in the developing embryo. Out of numerous considered annotated lncRNAs that we initially surveyed, our selection criteria led to a highly restricted set that contained epigenetically-regulated promoter signatures and cardiac-specific expression *in vivo*. Ablation of these transcripts in the developing mouse revealed modulatory roles of *Rubie, Atcayos, HrtLincR4, and HrtLincRX* within their local genomic environments. In particular, we found a requirement for the *Rubie* locus for normal *Bmp4* dosage.

Furthermore, loss of *Handlr*/*Hdn* and *Atcayos* led to minor reduction in LV mass in adult mice, along with modestly increased contractility among *Handlr*^-/-^ individuals. Loss of either of these two transcripts did not influence the hypertrophic or LV contractile response to pressure overload. Most importantly, despite clear transcription in the developing heart, none of the tested lincRNAs alone were required for embryo viability. However, when we generated compound heterozygotes for *Rubie* and *Bmp4*, we observed a slight yet consistent reduction in recovered offspring and a modest perturbation of right ventricular outflow tract orientation. Of course, this cohort of lincRNAs that we ablated does not include the entirety of all relevant cardiac transcripts. Nonetheless, we argue this set of transcripts represents a sufficient consideration of the early cardiac lincRNA landscape to predict modest individual roles for most noncoding transcripts of this type.

Similar to our observations, an independent study (Ritter, Ali et al. 2019) reported that the *Handlr/Hdn* locus produced multiple transcripts. Namely, *Hdn* was described as originating from the same TSS but splicing approximately 15kb 3’ beyond *Handlr,* along with *Hdnα*, which shares TSS and splice patterning (albeit 1 fewer exon) with *Handlr*. *Hdn/Hdnα* was described as localized to the cytoplasm near the nuclear envelope. Our results clearly supported *Handlr* enrichment in the nucleus, though we did not interrogate to which subnuclear component. Most importantly, they showed that complete deletion of the >20 kilobase *Hdn* locus produced embryonic-lethal cardiac and extraembryonic phenotypes. This accompanied perturbation of a cardiac gene expression program, including hyper-expression of *Hand2*. They attributed the act of transcription from the *Hdn*/*Hdnα* TSS as an important regulatory component of this phenotype, whereas *Hdn*(α)-specific RNA molecules were dispensable. In our work, ablation of the *Handlr/Hdn/Hdnα* TSS, which terminated downstream transcription, resulted in no lethality, dramatic cardiac phenotype, or altered *Hand2* expression. Therefore, both sets of experiments agree on the lack of requirement for *Handlr/Hdnα* as a functional lincRNA during cardiogenesis, but disagree on a link between *Handlr/Hdn/Hdnα* transcription and *Hand2* expression. Importantly, a clear regulatory mechanism/effect for transcription per se from the *Handlr/Hdn/Hdnα* TSS has yet to be established. We predict the large genomic deletion reported in Ritter *et al*. 2019 affects critical cardiac enhancer elements within the *Hdn* locus. Therefore, future work must more definitively assign a mechanistic role for RNA synthesis, splicing, and/or molecular function at this locus before attributing them to the mild or severe phenotypes observed by either study.

The subtle effects created by ablation of our collection of cardiac lincRNAs are consistent with the results most often obtained by others’ efforts to knockout developmentally-specific lincRNAs (Nakagawa, Naganuma et al. 2011, Nakagawa, Ip et al. 2012, Zhang, Arun et al. 2012, Sauvageau, Goff et al. 2013, Goff, Groff et al. 2015, Lai, Gong et al. 2015, Amandio, Necsulea et al. 2016, Goudarzi, Berg et al. 2019). In most instances where strong phenotypes have been observed, conflicting regulatory mechanisms, *in vivo* versus *in vitro* effects, lack of *in vivo* reproducibility, and/or conflated DNA-/RNA-/transcription-/splicing-based mechanisms prevent clean parsing of the underlying importance of lincRNA molecules. However, several lincRNAs in our cohort did seem to function within the nucleus to impact gene expression in their local environments, including *Rubie’s* influence on *Bmp4*. Consequently, future experiments are needed to dissect the physical mechanisms that underlie these effects. We speculate that the vast majority of singular lncRNAs fit as cogs into the multifaceted regulatory architecture of the nucleus, which individually can be compensated for during organogenesis *in vivo*. More so we hypothesize that overt phenotypic impacts in most lincRNA-centric experiments will be observed only after additional contextual molecular components are also manipulated. It may prove more beneficial to interpret most nuclear lincRNAs as collective epigenomic components analogous to histone/DNA modifications-with potential capabilities to alter the physical properties and orientation of the genome and recruit regulatory machinery, more so than as individual gene-like elements.

## Author Contributions

Project design and direction: B.G.B. and M.R.G. All experiments: M.R.G., except mouse echocardiography and surgery: Q.D., Y.H. under supervision of S.M.H, QRT-PCR: A.N., and interpretation of cardiac anatomy: I.S.K and K.R. Manuscript writing: M.R.G. and B.G.B with contribution from all authors.

## Acknowledgements

We thank Junli Zhang (Gladstone Transgenic Core) for pronuclear injection, Hazel Salunga for help with echocardiography, and the Gladstone Histology and Microscopy Core for embryo heart sectioning and histology. We are also grateful to Judy Morgan, Leslie Goodwin, and Laura Reinholdt (JAX) for mouse production.

## Funding

This work was funded by NIH/NHLBI (R01HL114948 and Bench to Bassinet Program UM1HL098179) to B.G.B., NIH/NHLBI HL127240 to S.M.H, and a Graduate Scholarship co-sponsored by the American Heart Association and the Lawrence J. and Florence A. DeGeorge Charitable Trust (ID: 15PRE24470159) to M.R.G. I.S.K. was funded by the Society of Pediatric Anesthesia, UCSF Research Allocation Program, the Hellman Family Fund, and the Department of Anesthesia at UCSF. This work was also supported by an NIH/NCRR grant (C06 RR018928) to the J. David Gladstone Institutes and by The Younger Family Fund to B.G.B.

## Competing Interests

B.G.B is a co-founder of and owns equity in Tenaya Therapeutics. M.R.G. is an employee of and owns equity in Vascugen Inc. S.M.H. is an executive and shareholder of Amgen, Inc. and a co-founder with equity stake in Tenaya Therapeutics. These interests are not related to the work described here.

## Materials and Methods

### Informatic search for cardiac lincRNAs

Raw stranded, total RNA-seq reads from ESC (day 0), MES (day 4), CP (day 5.3), and CM (day 10) stages (Wamstad, Alexander et al. 2012) and cMES (day 4.75) stage (Devine, Wythe et al. 2014) of mESC *in vitro* differentiation into cardiomyocytes were mapped to the mouse genome (mm9) and aligned to Noncode v4 (Xie, Yuan et al. 2014) annotated lncRNAs using the Cufflinks suite of software (Trapnell, Williams et al. 2010). ChIP-seq domains positive for trimethylation of histone 3 lysine 4 (H3K4me3), acetylation of histone 3 lysine 27 (H3K27Ac), and trimethylation of histone 3 lysine 27 (H3K27me3) for ESC, MES, CP, and CM stages were obtained from Wamstad, Alexander et al. The following criteria were used to generate a candidate list of lincRNAs. 1.) Less than 0.5 fragments per kilobase per million reads (FPKM) in mESCs. 2.) Greater than 1.0 FPKM at CP or cMES stage of differentiation (expression at other time points was not factored into selection). 3.) Positive H3K4me3 ChIP-seq signal at TSS during CP stage. 4.) Positive H3K27Ac at TSS during CP stage. 5.) At least 1 exon splice in transcript. 6.) No splice events into neighboring protein coding genes 7.) TSS at least 1kb from nearest protein-coding gene TSS. Screened candidates tracks were then visually inspected via UCSC Genome Browser (Kent 2002) to filter for lincRNAs with expression patterns that matched the assigned lincRNA structure and not simply spurious reads at the general locus.

### Analysis of lincRNA coding potential

PhyloCSF (Lin, Jungreis et al. 2011) browser tracks were uploaded to the UCSC Genome Browser for interrogation. Ficket and hexamer scores were calculated with CPAT software (Wang, Park et al. 2013) and visualized with the ‘ggplot2’ package (Wickham 2016) in R version 3.4.0 (R-Core-Team 2017). CNCI scores were obtained from NONCODEv4 annotations (Xie, Yuan et al. 2014).

### mESC differentiation and cardiac progenitor cell fractionation

Directed cardiomyocyte differentiations were performed as previously described (Wamstad, Alexander et al. 2012) using the *Smarcd3-F6nlsEGFP* mESC line (Devine, Wythe et al. 2014) with minor modifications to improve differentiation efficiency. Briefly, three days before differentiation induction (day -3), mESCs were split into 2i + LIF media on gelatin. The following day (day -2), 2i + LIF was replaced with 15% FBS (HiClone) in DMEM + 1X non-essential amino acids + 1X sodium pyruvate + 1X GlutaMAX + 1X βmercaptoethanol + 1X penicillin/streptomycin + 1000U/ml LIF (ESGRO, EMD). The following day (day -1), cells were fed again with the same 15% FBS-LIF media to complete their conversion to epiblast-like stem cells. One day later (day 0), cardiac differentiation was initiated as per Wamstad, Alexander et al. On day 5.3, CP cells were dissociated with TrypLE (Gibco) and quenched in DMEM:F12 (Gibco) + 10% FBS (HiClone). After washing in D-PBS, nuclei were isolated using the Nuclei EZ Prep kit (SigmaAldrich), pelleted, and supernatant was collected as cytoplasmic fraction. Nuclei were washed again, and pellet was harvested with Trizol (Invitrogen) as nuclear fraction in subsequent qPCR quantification experiments. Cytoplasmic fractions were processed equivalently.

### Whole mount in situ hybridization

Primers were designed to amplify *in situ* probe templates between 440bp and 1.5kb for each candidate lincRNA off cDNA from CP stage of *in vitro* differentiation (Supplemental table S2). Templates were electrophoresed in 1.0% agarose gel and purified using QIAquick gel extraction kit (Qiagen). These templates were then TOPO TA cloned into pCR4-TOPO using the TOPO TA cloning kit (Invitrogen) and Sanger sequenced to validate orientation in plasmid and proper composition. 2μg linearized vector for each lincRNA template were then input into digoxygenin (DIG) RNA synthesis kit reactions (Roche) in 40μL total volume using either T7 or T3 primers, depending on template orientation. Transcription was carried out for 2 hours at 37°C. Afterward, 8U DNase I (NEB) were added to each reaction and incubated for 15 min at 37°C to degrade DNA. DNase reactions were quenched with 1.5μL EDTA, and DIG-RNA probes were cleaned and concentrated with RNeasy Mini Columns (Qiagen), EtOH precipitated and washed, and resuspended in 20μL H_2_O. DIG probes were then diluted to 100μg/mL in HYB buffer (50% formamide + 5X SSC pH 4.5 + 50μg/mL yeast tRNA + 75μg/mL heparin + 0.2% Tween-20 + 0.5% CHAPS + 5mM EDTA). E7.5 through E12.5, mouse embryos were liberated from the uterus and dissected from extraembryonic tissues and membranes. Embryos were washed with D-PBS and fixed overnight in 4% paraformaldehyde and then washed 3x in PBT (PBS + 0.1% Tween-20) on ice. Embryos were dehydrated in MeOH series (25%, 50%, 75%, 2x 100%, 5 min each). Then, samples were rehydrated by reversing this series including 2 extra PBT washes. Embryos were bleached in 6% H_2_O_2_ in PBT for 15 minutes at RT with rocking. Embryos were washed 3 x 5 min in PBT and treated with 10ug/mL proteinase K for 5 min (E7.5), 10 min (E8.5), 20 min (E9.5), or 30 min (E10.5+) rocking at RT and then quenched 2x with 2mg/mL glycine in PBT followed by 3 x 5min washes in PBTw. Embryos were re-fixed in 4% paraformaldehyde + 0.2% glutaraldehyde for 20 min with rocking and washed an additional 5x 5min with PBT. Embryos were then rinsed 2x in 65°C HYB buffer and incubated in HYB buffer for 3 hours at 65°C. Then, lincRNA-specific probes (in HYB) were added, respectively, to final concentration of 1μg/mL and hybridized overnight at 65°C. Embryos were rinsed 3 x 5 min in 65°C WASH1 buffer (50% formamide + 5X SSC pH 4.5 + 1% SDS) and then incubated 2 x 30 min again in 65°C WASH1 buffer. Next, embryos were washed 2 x 30 min in 65°C WASH2 (50% formamide + 2X SSC pH 4.5 + 0.1% Tween-20), followed by 3 x 5 min RT washes in TTBS (25mM Tris HCl pH 7.4 + 135mM NaCl + 2.5mM KCl + 0.1% Tween-20). Embryos were then blocked in TTBS containing 20% sheep serum for 3 hours at RT and stained overnight with alkaline phosphatase (AP) conjugated anti-DIG Fab fragments in TTBS + 1% sheep serum (1:5000, Roche). Embryos were then rinsed 3x 5min in RT TTBS, followed by 6x 1hr TTBS washes at RT. A final TTBS wash was then performed overnight at 4°C. Embryos were then washed 2 x 30 min in AP buffer (100mM Tris pH 9.5 + 50mM MgCl_2_ + 100mM NaCl + 0.1% Tween-20) at RT. Then, Boehringer Purple AP substrate was added to embryos to initiate staining reactions. Reactions were allowed to progress in the dark until suitable contrast was observed. AP reactions were quenched with 3x PBT washes containing 1mM EDTA, followed by multiple PBT pH5.5 washes. A final fixation was then performed overnight in 4% paraformaldehyde and 0.1% glutaraldehyde at 4°C. Finally, embryos were dehydrated again in methanol series and stored in 100% MeOH at -20°C. Embryos were imaged on an upright microscope, and images were white balanced with Adobe Photoshop.

### Cas9 lincRNA knockout, mouse husbandry, and genotyping

All mouse experiments were carried out in accordance with IACUC protocols and cared for by the UCSF LARC. For each lincRNA, two cut sites were targeted to induce a 2-3kb deletion flanking the TSS/promoter. Two sequence-specific truncated single guide RNA (tru-gRNA, Supplemental table S3) (Fu, Sander et al. 2014) regions were separately cloned into pX330 (Addgene). After generating T7 promoter-containing sgRNA templates by PCR using Phusion TAC polymerase (NEB), tru-sgRNAs were transcribed using the Hiscribe T7 High Yield RNA Synthesis Kit (NEB). Tru-sgRNA was extracted using Trizol reagent (Invitrogen) and dual chloroform purifications before immunoprecipitating with isopropanol. Each tru-gRNA pair was then resuspended in sterile 5mM Tris-HCl before pronuclear injection by the Gladstone transgenic mouse core. Injections were carried out as previously described (Yang, Wang et al. 2014). To increase efficiency of obtaining deletions for each target site, all pairs were co-injected into each of 70 FVB/n pronuclei. All genotyping was performed on tail clips stored at - 20°C. To extract gDNA, tail clips were suspended in 100μL 50mM NaOH in H_2_O and incubated at 95°C for 40 minutes. Tubes were agitated to break up tissue, and remaining solids were allowed to settle before use. pH was normalized by addition of 7.0μL of 1M Tris HCl pH 7.4. 1.5μL was then input into PCR reactions using Q5 2X master mix (NEB) and 3 gene-specific primers for simultaneous WT and KO product amplification (Supplemental table S4). Reactions were carried out according to manufacturer-specified recommendations. F0 founders for single lincRNA deletions were first identified and bred into C57BL/6j to establish germline transmission (F1). Separate F1 heterozygotes for each individual lincRNA deletion were then outbred into the C57BL/6j background for multiple generations to reduce off-target effects. *Handlr* and *Atcayos* null alleles were generated in collaboration with Jackson Laboratory using the same targeting strategy but in a homogenous C57BL/6j background.

### Transverse aortic constriction cardiac hypertrophy models

Surgery was performed under IACUC protocols and monitored by the UCSF LARC. Experiments were performed as described (Duan, McMahon et al. 2017). For transverse aortic constriction (TAC), 12-20 week-old male mice were anaesthetized with ketamine/xylazine and mechanically ventilated. After thoracotomy, TAC was executed between the left common carotid and the brachiocephalic arteries using a 7-0 silk suture and 27-gauge needle. After surgery, pressure overload was confirmed by Doppler probe measurement of flow velocity at the carotid artery. Echocardiography was performed at baseline, 1 week, 4 weeks, 6 weeks, and 8 weeks after operation to measure left ventricle (LV) fractional area change (%FAC). LV areas were obtained from two-dimensional measurements at the end-diastole and end-systole. At baseline, non-biological echocardiography variability required outlier removal. Week 0 (pre-TAC) mice measured to be in cardiac failure (%FAC < 30.0%) with increased %FAC week 1 after TAC were deemed failed measurements. 1-2 instances of this were observed in all groups. From remaining data, ‘1.5X interquartile range rule’ was used to eliminate outliers. The LV mass was estimated by M-mode measurements and the equation *M_LV_* = ((*IVS_D_* + *LVID_D_* + *LVPW_D_*)^3^ − *LVID_D_*^3^) ×1.053 (Marwick, Gillebert et al. 2015) M_LV_, left ventricular mass; IVS_D_, diastolic interventricular septum width; LVID_D_, diastolic left ventricular internal diameter; LVPW_D_, diastolic left ventricular posterior wall thickness.

At 8 weeks post-surgery, mice were sacrificed for analysis. First, left ventricle, lung, and body weights were measured. Subsequently, a 10-20mg concentric short axis slice of the left ventricle was collected and preserved in RNAlater reagent (ThermoFisher). Heart sections were disrupted in PureZOL (Bio-Rad) on a TissueLyser II (Qiagen). RNA was then purified with Aurum purification kit (BioRad). qRT-PCR was performed using TaqMan chemistry including FastStart Universal Probe Master (Roche), labeled probes from the Universal Probe Library (Roche), and gene-specific oligonucleotide primers run on a 7900HT (ThermoFisher) cycler with absolute quantification. Gene expression levels were normalized to *cycloB* and *Actb* internal controls using the *Δ*Ct method.

### E8.25 RNA isolation and qPCR analysis

At E8.25, embryos were removed from the uterus and dissected from extraembryonic tissues and membranes. Only embryos displaying late cardiac crescent formation before heart tube expansion and cavitation were kept and deemed to be at E8.25. The anterior half of each embryo was washed twice in cold PBS and transferred into Trizol (Invitrogen), while the posterior half was washed in PBS and stored at -20°C for genotyping. RNA from Trizol samples was precipitated using standard protocols and further purified/ condensed using Qiagen RNeasy MinElute columns. 250ng RNA was reverse transcribed using the AffinityScript Reverse Transcription kit (Agilent) using 200ng random hexamer and/or 100ng dT_20_ primers, where appropriate. RT-qPCR was subsequently performed with 5.0ng cDNA and 500nM gene-specific primers (Supplemental table S1) in PowerUP SYBR Green master mix (Thermo Fisher). Reactions were run on a 7900HT (ThermoFisher) cycler with absolute quantification. Gene expression levels were normalized to *Actb* internal controls using the *Δ*Ct method.

## E15.5 Histology

At E15.5, embryos were liberated from the uterus and dissected from extraembryonic tissues and membranes. Whole hearts were removed, rinsed twice in D-PBS, and fixed overnight in 4% paraformaldehyde. Each heart was then paraffin embedded and sectioned at an oblique transverse plane for four chamber visualization. Hematoxylin and eosin staining and imaging were performed by the Gladstone Histology Core (UCSF).

---

**Supplemental Figure S1.**
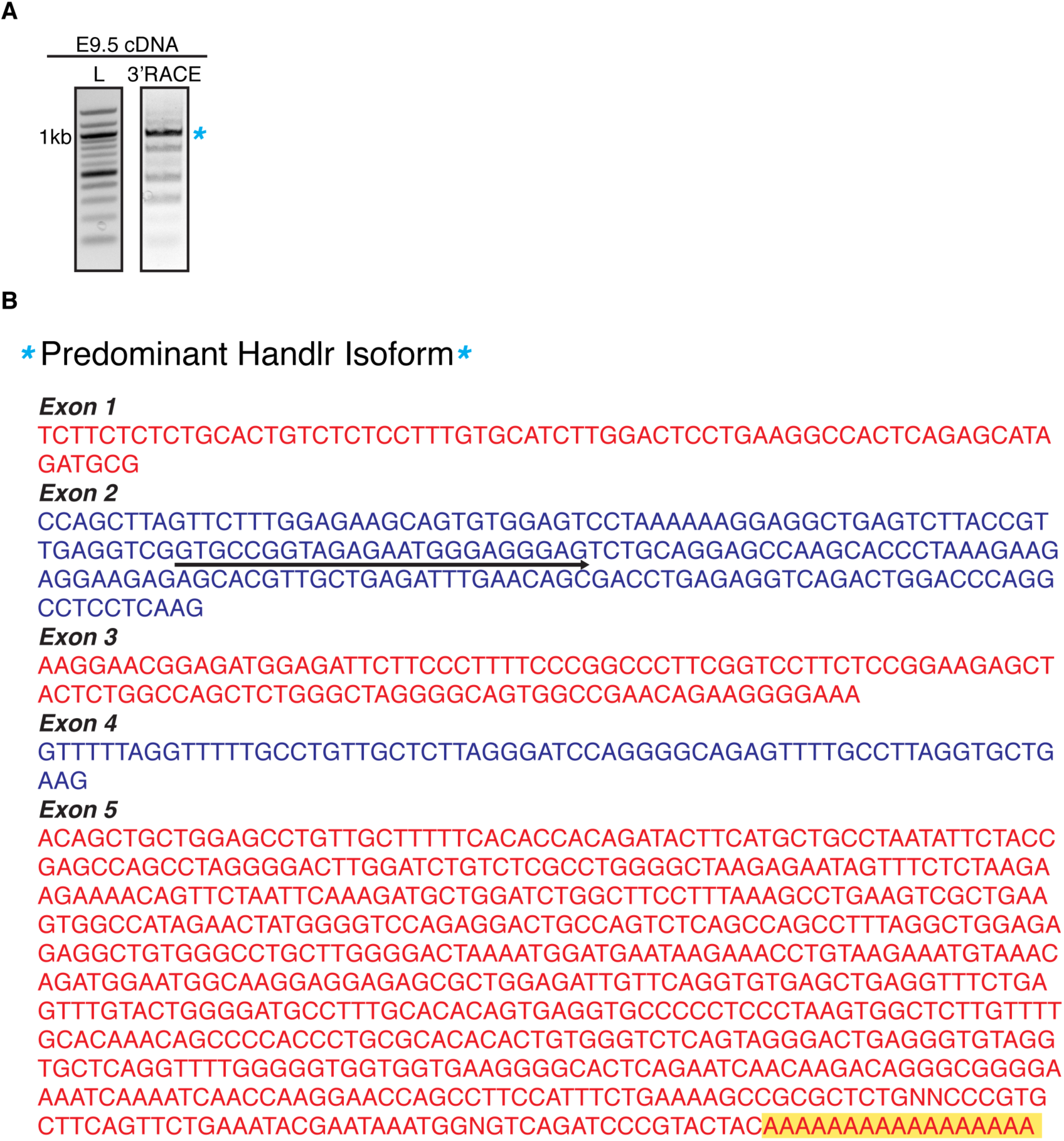
Predominant *Handlr* transcript *in vivo*. A. Electrophoresed Handlr 3’ RACE PCR products from E9.5 mouse cDNA. B. Sanger sequence of predominant RNA transcript, polyA tail highlighted; T in place of U due to DNA sequencing; arrow, location of 3’RACE *Handlr*-specific primer.

**Supplemental Figure S2.**
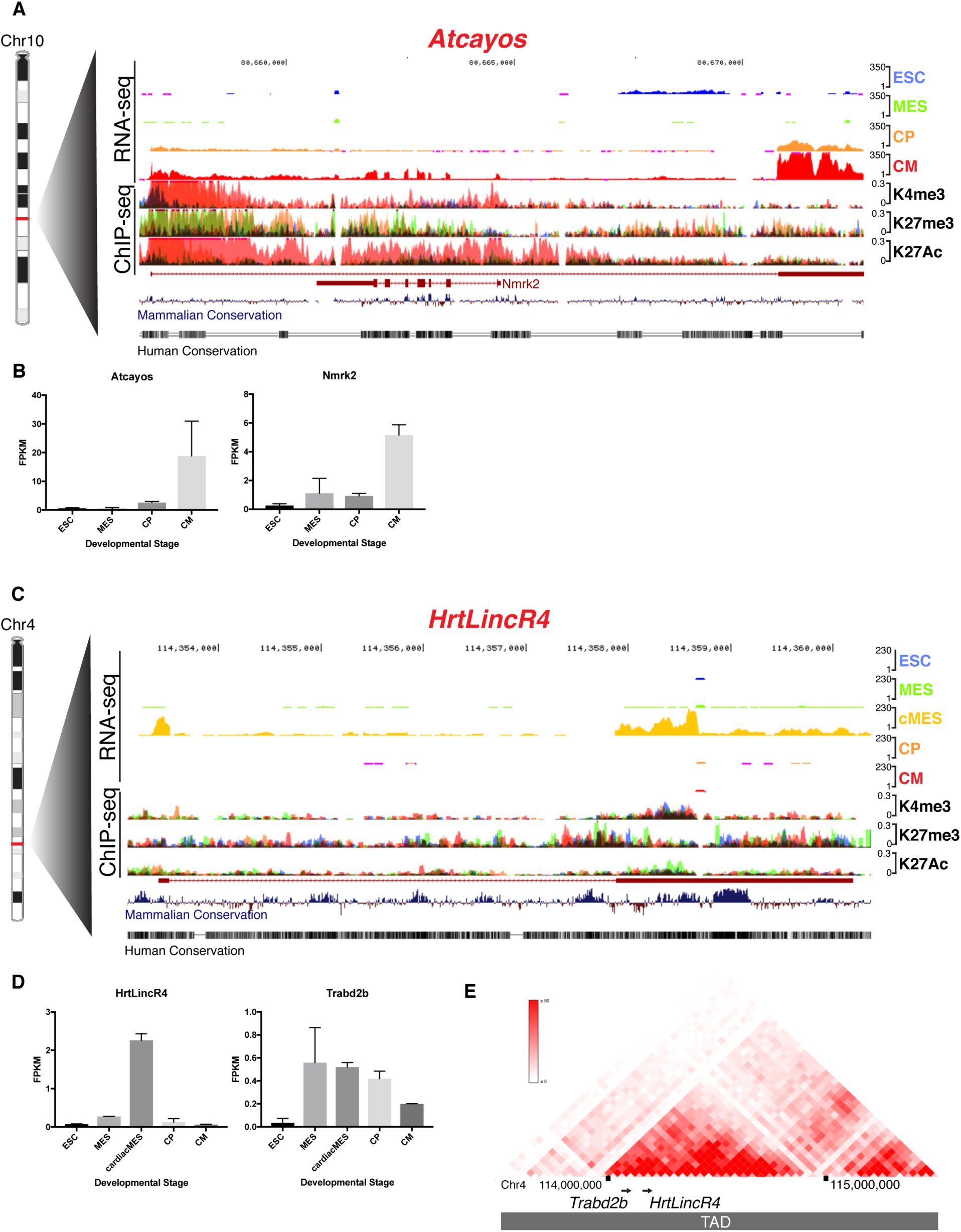
Genomic characterization of *Atcayos* and *HrtLincR4 in vitro*. A. UCSC Genome Browser tracks of *Atcayos* and *Nmrk2* RNA-seq and overlaid histone H3 ChIP-seq at ESC, MES, CP, and CM stages of *in vitro* differentiation. B. Quantified expression of Atcayos and Nmrk2 at each differentiation stage. C. UCSC Genome Browser tracks of HrtLincR4 RNA-seq and overlaid histone H3 ChIP-seq during cardiac differentiation *in vitro*; D. Quantified expression of *HrtlincR4* and *Trabd2b* at each differentiation stage. E. 3D Genome Browser Hi-C heatmap of chromosome interactions around *HrtlincR4* and *Trabd2b* loci. ESC, embryonic stem cell, MES, mesoderm, CP, cardiac progenitor, CM, cardiomyocyte; blue, ESC, green, MES, orange, CP, red, CM; K4me3, histone H3 lysine 4 trimethylation; K27me3, histone H3 lysine 27 trimethylation; K27Ac, histone H3 lysine 27 acetylation; TAD, topologically associated domain. Ensembl annotations in red.

**Supplemental Figure S3.**
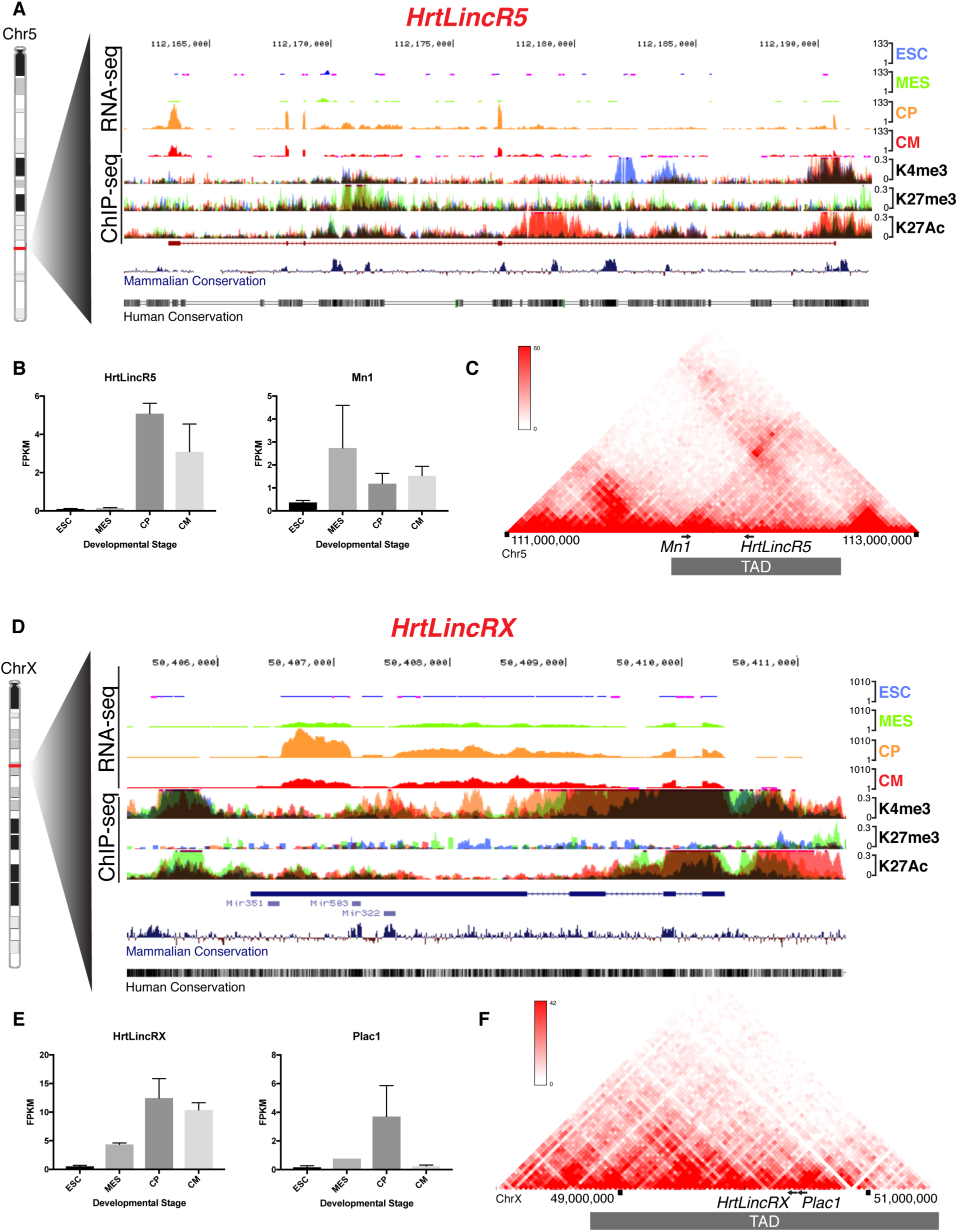
Genomic characterization of *HrtLincR5* and *HrtLincRX in vitro*. A. UCSC Genome Browser tracks of *HrtLincR5* RNA-seq and overlaid histone H3 ChIP-seq during cardiac differentiation *in vitro*. Ensembl annotation in red. B. Quantified expression of HrtlincR5 and Mn1 at each differentiation stage. C. 3D Genome Browser Hi-C heatmap (of chromosome interactions around *HrtlincR5* and *Mn1* loci. D. Genome Browser tracks of *HrtLincRX* RNA-seq and overlaid histone H3 ChIP-seq during cardiac differentiation *in vitro*; E. Quantified expression of *HrtLincRX* and *Plac1* at each differentiation stage. F. 3D Genome Browser Hi-C heatmap of chromosome interactions around *HrtlincRX* and *Plac1* loci. ESC, embryonic stem cell, MES, mesoderm, cMES, cardiac mesoderm; CP, cardiac progenitor, CM, cardiomyocyte; blue, ESC; green, MES; orange, CP; red, CM; K4me3, histone H3 lysine 4 trimethylation; K27me3, histone H3 lysine 27 trimethylation; K27Ac, histone H3 lysine 27 acetylation; RefSeq annotation in blue. RefSeq annotations, including 3’ miRNA cluster, in blue.

**Supplemental Figure S4.**
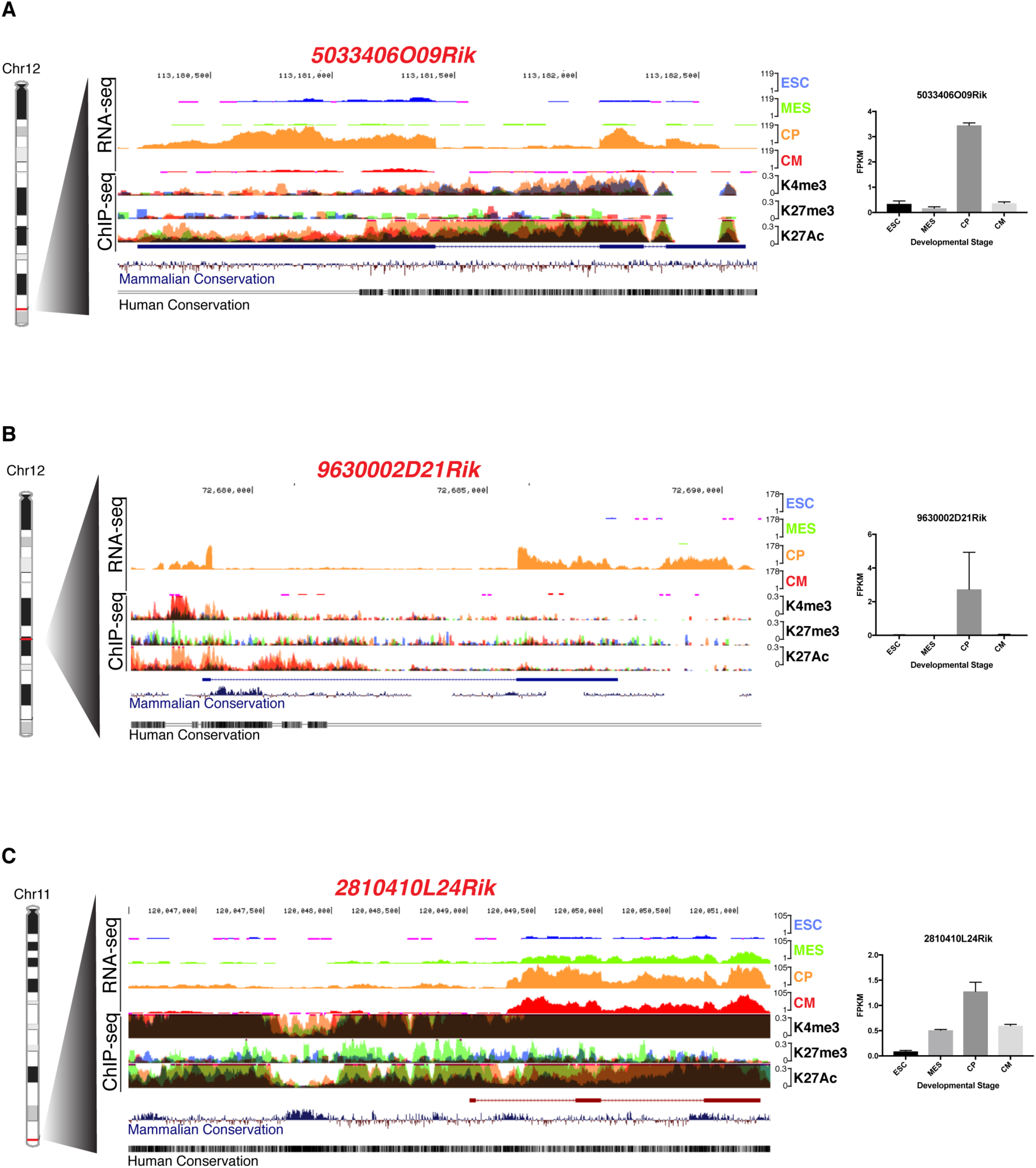
Genomic characterization of *5033406O09Rik*, *9630002D21Rik*, and *2810410L24Rik in vitro*. A. UCSC Genome Browser tracks of *5033406O09Rik* RNA-seq and overlaid histone H3 ChIP-seq during cardiac differentiation *in vitro*, as well as quantified expression at each differentiation stage. B. UCSC Genome Browser tracks of *9630002D21Rik* RNA-seq and overlaid histone H3 ChIP-seq during cardiac differentiation *in vitro*, as well as quantified expression at each differentiation stage. C. UCSC Genome Browser tracks of *2810410L24Rik* RNA-seq and overlaid histone H3 ChIP-seq during cardiac differentiation *in vitro*, as well as quantified expression at each differentiation stage. ESC, embryonic stem cell, MES, mesoderm, cMES, cardiac mesoderm; CP, cardiac progenitor, CM, cardiomyocyte; blue, ESC; green, MES; orange, CP; red, CM; K4me3, histone H3 lysine 4 trimethylation; K27me3, histone H3 lysine 27 trimethylation; K27Ac, histone H3 lysine 27 acetylation; Ensembl annotation in red, RefSeq annotations in blue.

**Supplemental Figure S5.**
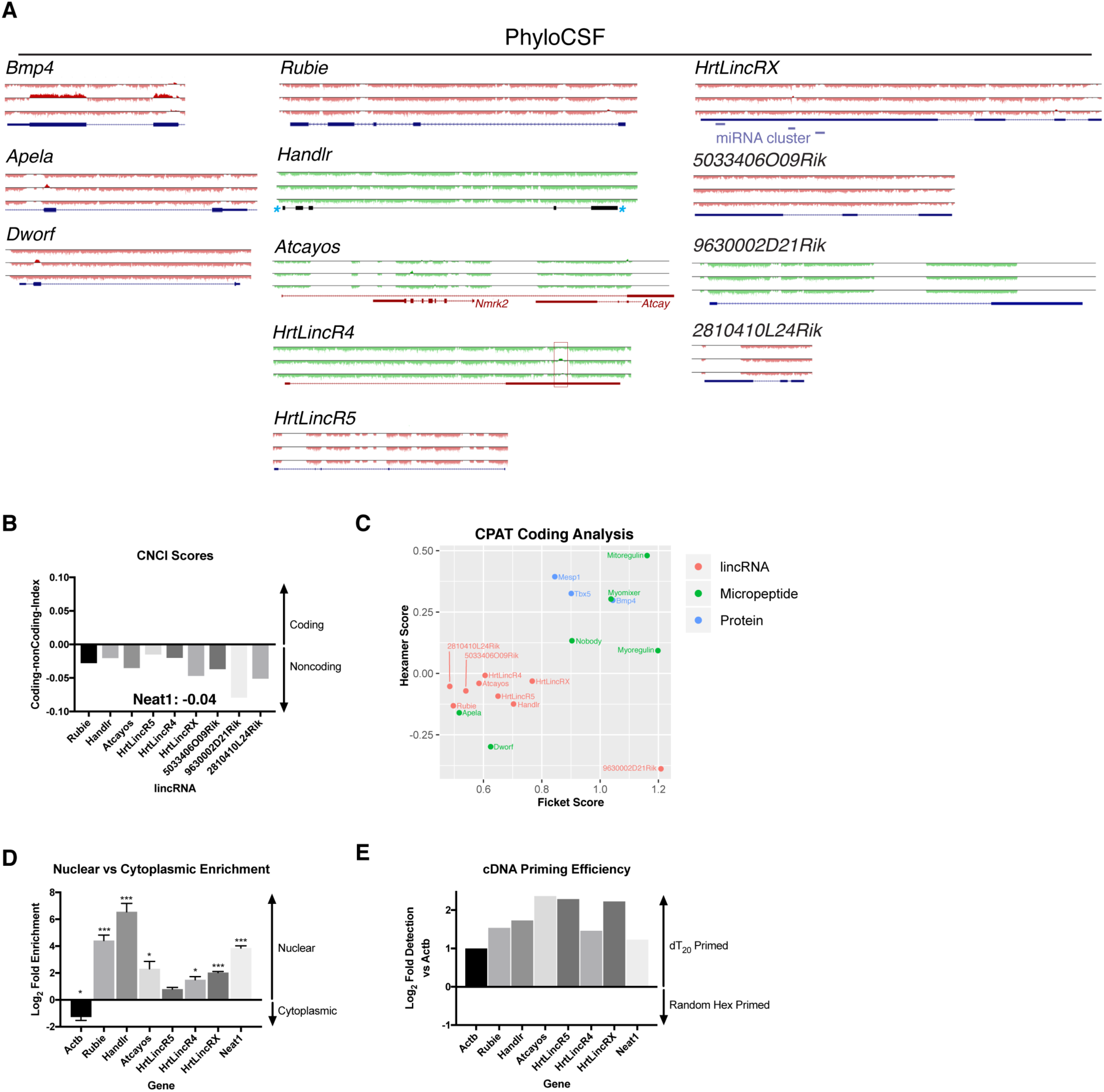
Molecular characterization of lincRNA cohort. A. UCSC Genome Browser tracks of PhyloCSF codon scores for all three frames of known protein coding gene *Bmp4*, micropeptide coding genes *Apela* and *Dworf*, and lincRNA cohort; red box, potential 28 amino acid-coding open reading frame in *HrtLincR4*; scale, -15 to +15; positive score indicates higher coding potential; green, (+) strand; red, (-) strand. B. Coding-non-Coding-Index (CNCI) scores for lincRNA cohort compared to known lincRNA Neat1. C. Comparison of CPAT algorithm calculations of Ficket and Hexamer scores for known protein coding genes, micropeptide coding genes, and our lincRNA cohort. D. Nuclear vs cytoplasmic enrichment of lincRNA cohort compared to Actb and known nuclear-enriched lincRNA Neat1; *, p < 0.05; ***, p < 0.005; n.s., not significant; Student’s 2-tailed t-test. E. Efficiency of RT-qPCR amplification from dT_20_- or random hexamer-primed cDNA for lincRNA cohort compared to *Actb* and known polyadenylated lincRNA *Neat1*. Data presented as mean +/-SEM.

**Supplemental Figure S6.**
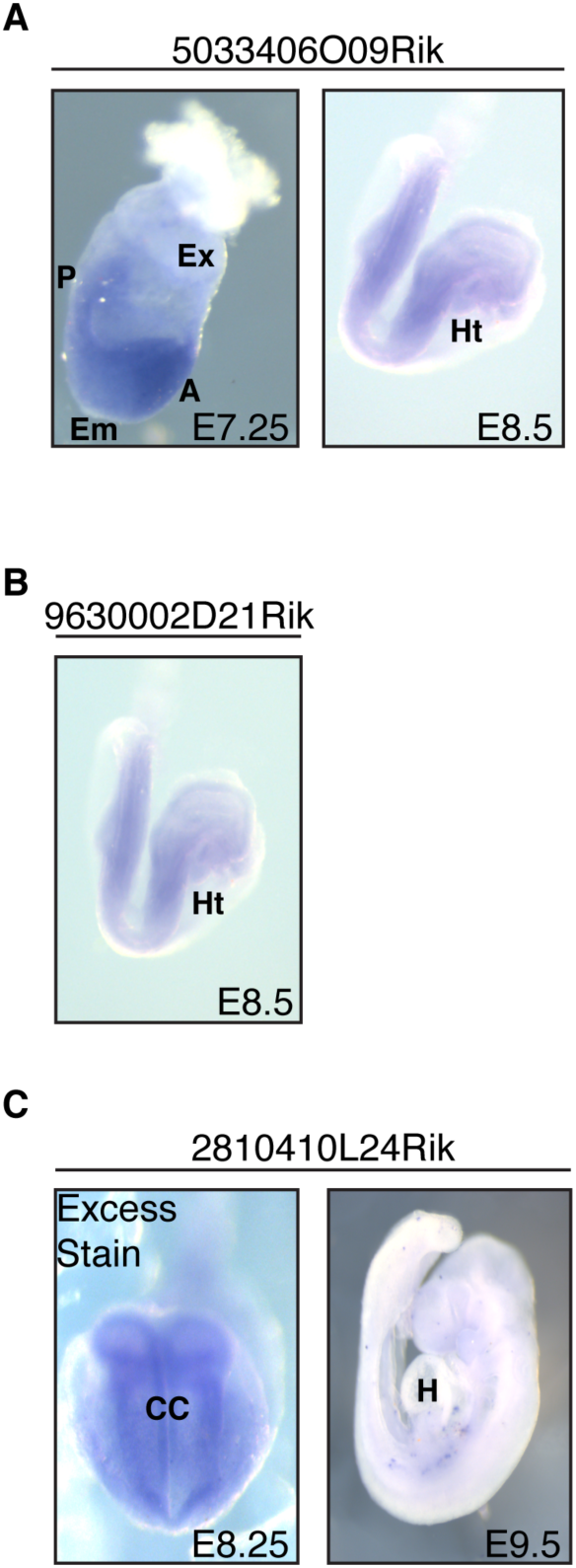
lincRNA expression patterns *in vivo*. A. *in situ* hybridization staining for *5033406O09Rik* at E7.25 and E7.5. B. *in situ* hybridization staining for *9630002D21Rik* at E8.5. C. *in situ* hybridization staining for *2810410L24Rik* at E8.25 and E9.5. A, anterior; Em, embryonic region; Ex, extraembryonic region; P, posterior; CC, cardiac crescent; Ht, heart tube; H, heart.

**Supplemental Figure S7.**
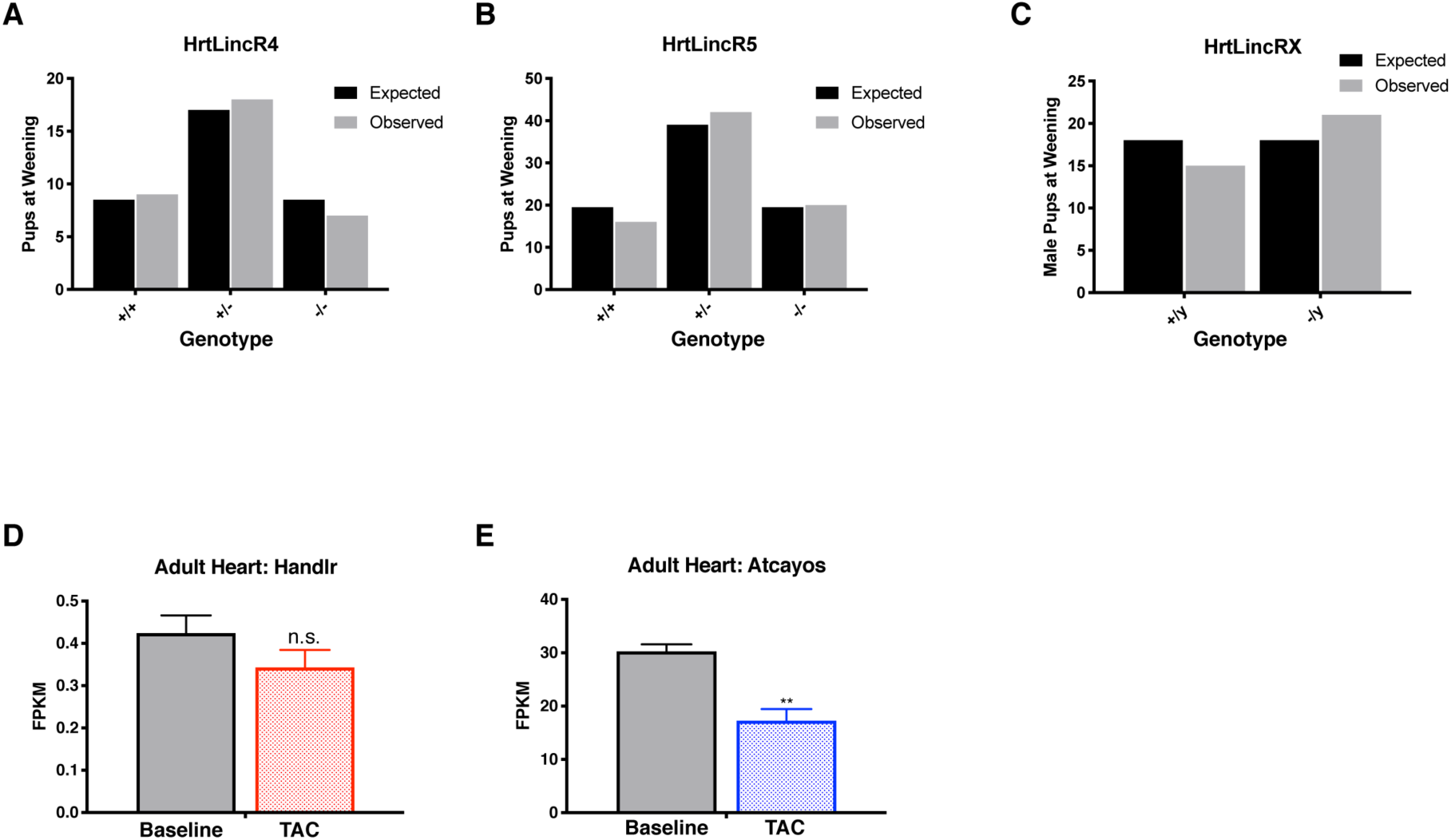
Viability and phenotypic effects after lincRNA knockout. A. Offspring recovered at weaning from *HrtLincR4^+/-^*x *HrtLincR4^+/-^* cross vs expected Mendelian ratios. B. Offspring recovered at weaning from *HrtLincR5^+/-^* x *HrtLincR5^+/-^*cross vs expected Mendelian ratios. C. Male offspring recovered at weaning from *HrtLincRX^+/-^* x *HrtLincRX^+/y^* cross vs expected Mendelian ratios. D. RNA-seq expression of Handlr in adult heart before and after TAC. E. RNA-seq expression of Atcayos in adult heart before and after TAC from Duan et al, 2017. **, p < 0.01, Student’s 2-tailed t-test; TAC, transverse aortic constriction; WT, wild type.

**Supplemental Figure S8.**
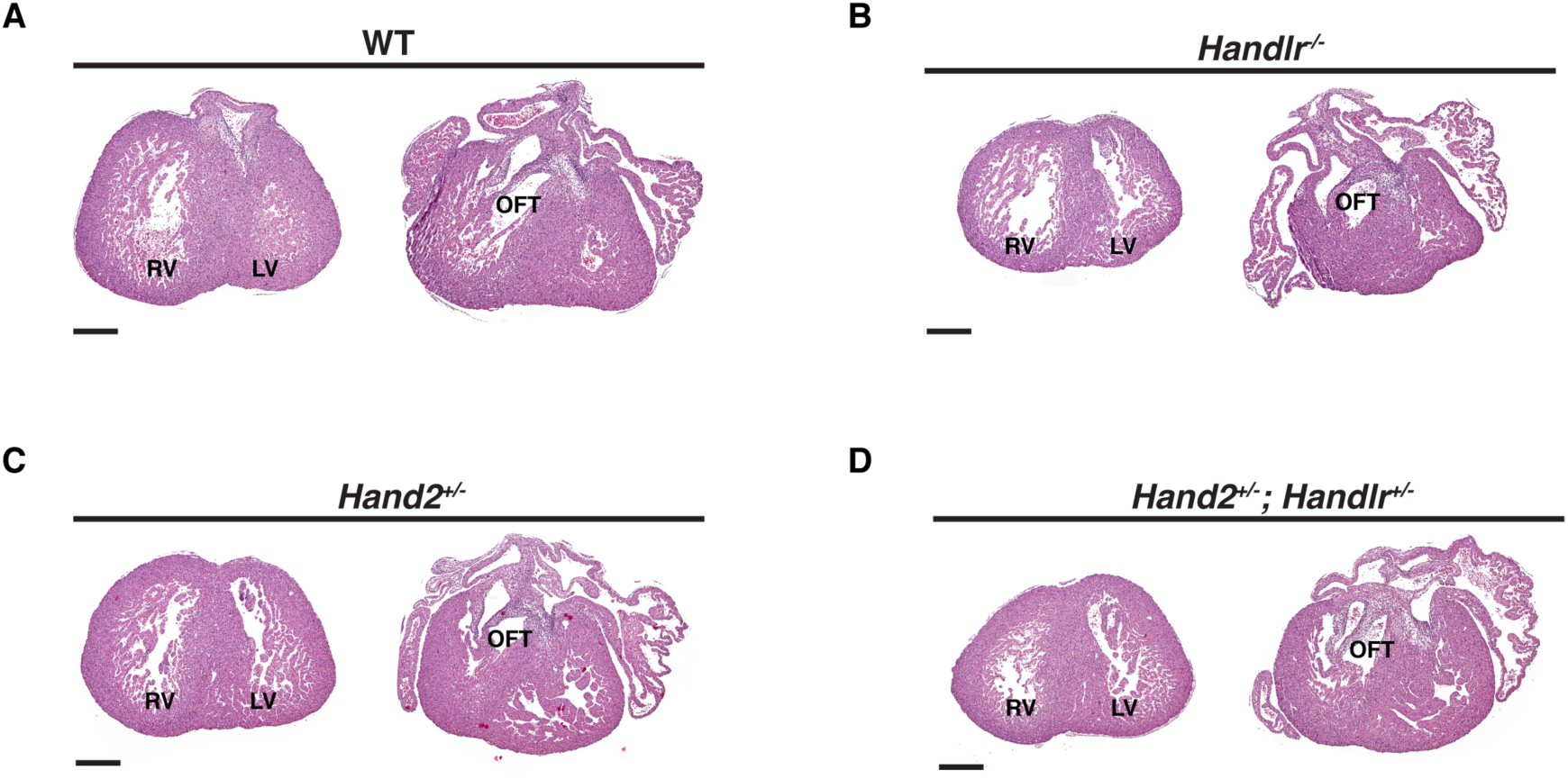
Effect of *Handlr* ablation on heart development at E15.5. A-D. Oblique transverse hematoxylin and eosin histological sections of cardiac ventricular and OFT morphogenesis, respectively, at E15.5. A. Representative WT morphology. Representative *Handlr^-/-^*morphology. C. Representative *Hand2^+/-^* morphology. D. Representative *Hand2^+/-^; Handlr^+/-^* morphology. RV, right ventricle; LV, left ventricle; OFT, outflow tract; WT, wild type; scale bar, 300μm.

**Supplemental Table S1.**
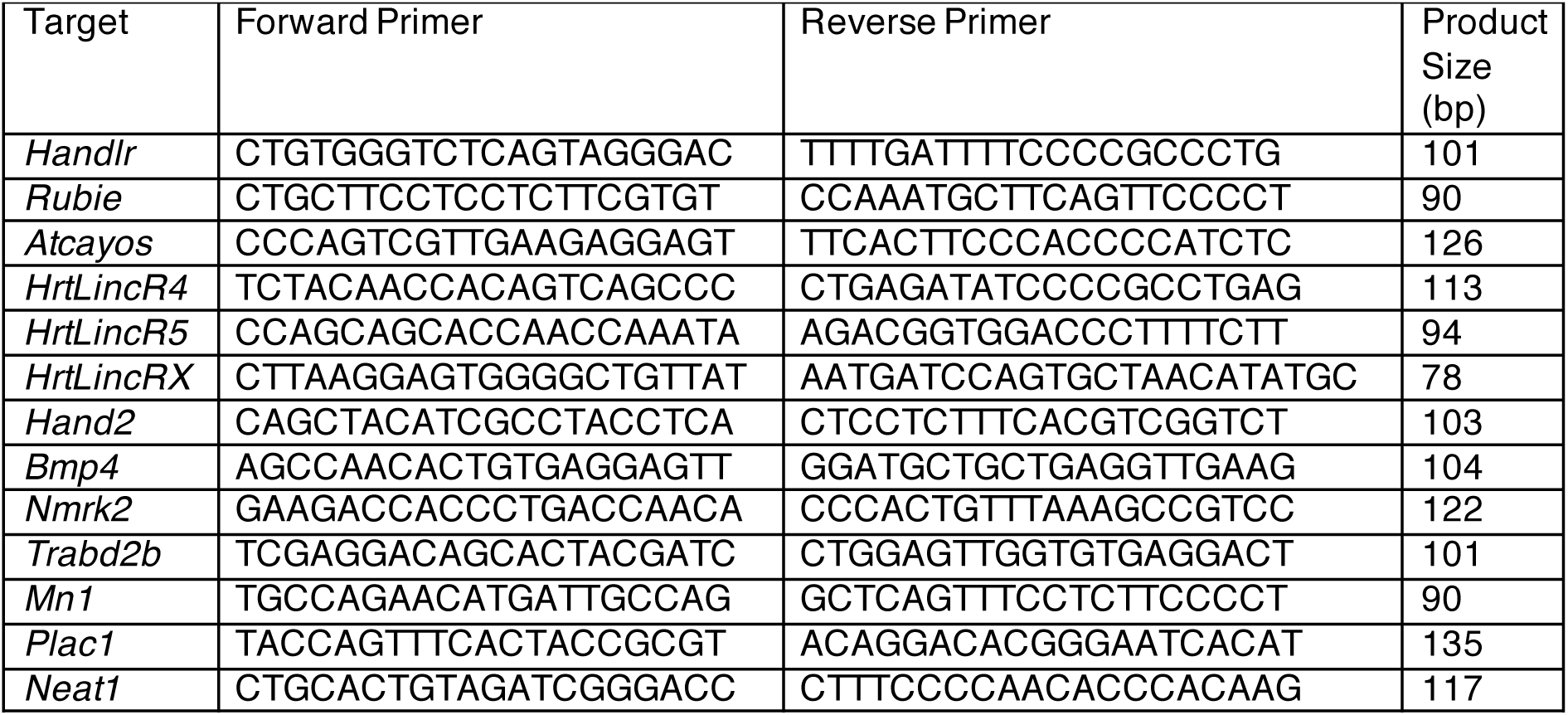
RT-qPCR primers

**Supplemental Table S2.**
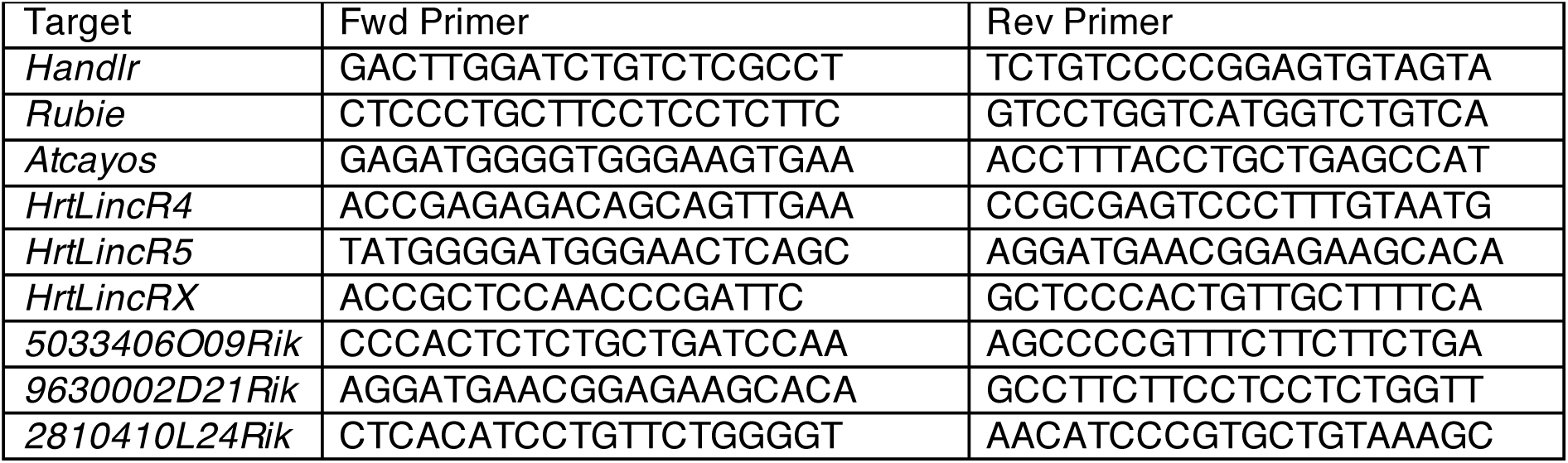
*in situ* hybridization probe primers

**Supplemental Table S3.**
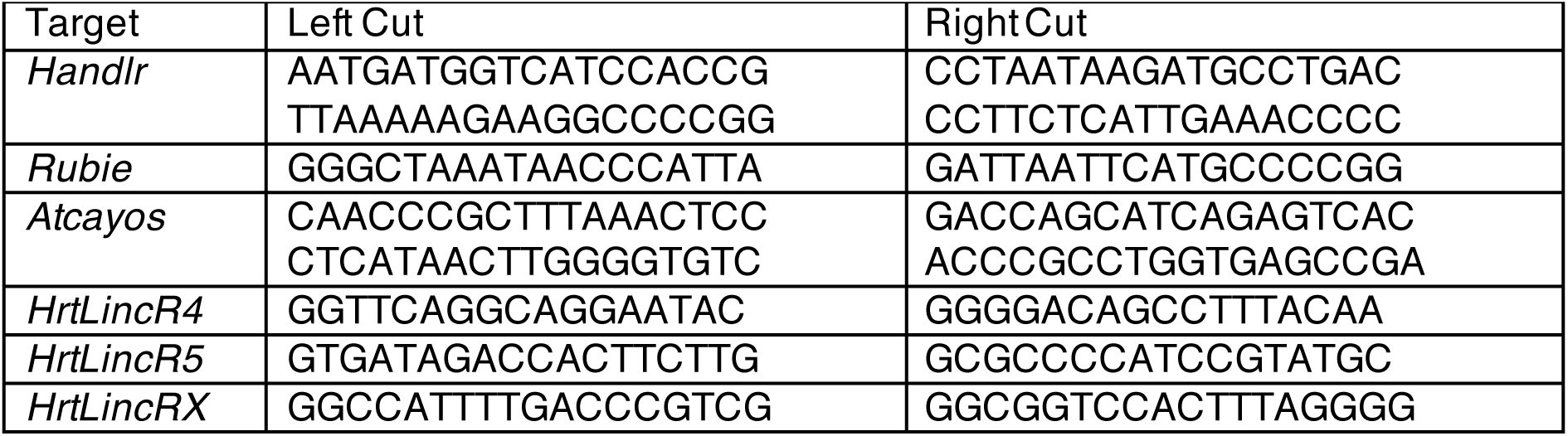
tru-sgRNA oligomers

**Supplemental Table S4.**
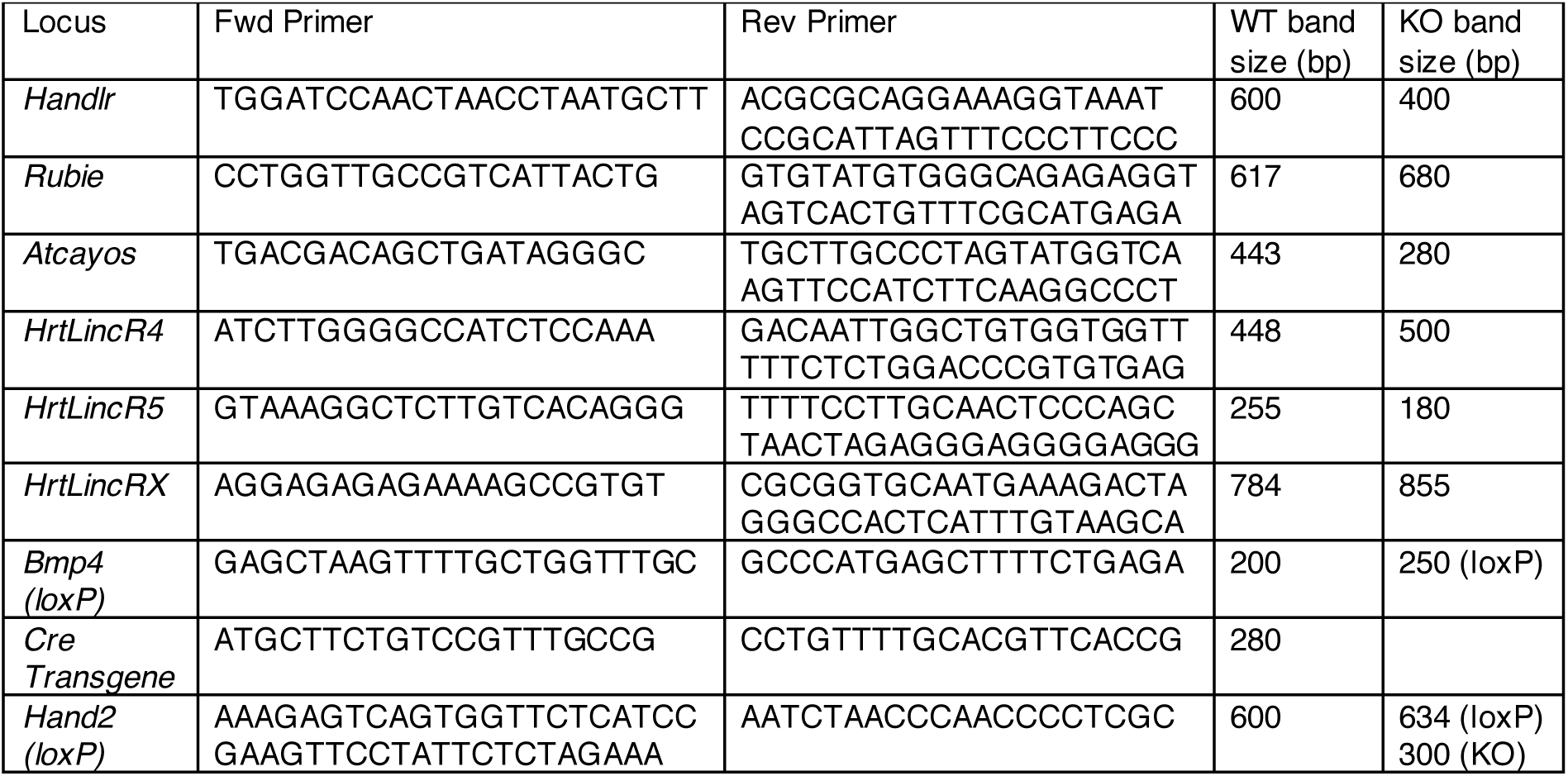
Genotyping primers

**Supplemental Table S5.** *Handlr* KO TAC echocardiography

| Time Point | *Handlr*<sup>+/+</sup> |  |  |  |  | *Handlr*<sup>-/-</sup> |  |  |  |  |
| --- | --- | --- | --- | --- | --- | --- | --- | --- | --- | --- |
|  | Baseline | Week 1 | Week 4 | Week 6 | Week 8 | Baseline | Week 1 | Week 4 | Week 6 | Week 8 |
| ENDOarea;d (mm<sup>2</sup>) | 16.21±0.71 | 16.26±1.19 | 16.65±1.42 | 16.19±1.58 | 17.35±1.63 | 14.00±0.70 | 14.63±0.29 | 15.39±1.16 | 15.65±0.49 | 16.05±1.09 |
| ENDOarea;s (mm<sup>2</sup>) | 10.07±0.45 | 11.82±1.15 | 12.59±1.10 | 12.30±1.17 | 13.35±1.36 | 7.51±0.53 | 9.72±0.35 | 10.68±1.25 | 11.38±0.27 | 12.45±1.04 |
| IVS;d (mm) | 0.64±0.04 | 1.02±0.05 | 0.90±0.06 | 1.12±0.06 | 1.20±0.08 | 0.63±0.04 | 0.93±0.04 | 1.13±0.04 | 1.05±0.07 | 1.29±0.06 |
| IVS;s (mm) | 0.96±0.04 | 1.38±0.04 | 1.16±0.08 | 1.25±0.05 | 1.39±0.05 | 0.91±0.02 | 1.28±0.07 | 1.33±0.06 | 1.28±0.05 | 1.46±0.11 |
| LVID;d (mm) | 4.54±0.10 | 4.38±0.16 | 4.65±0.19 | 4.58±0.24 | 4.68±0.19 | 4.19±0.09 | 4.25±0.04 | 4.49±0.18 | 4.57±0.06 | 4.43±0.13 |
| LVPW;d (mm) | 0.66±0.05 | 1.00±0.05 | 1.03±0.11 | 1.22±0.07 | 1.13±0.04 | 0.61±0.04 | 0.89±0.09 | 0.93±0.09 | 1.03±0.05 | 1.11±0.05 |
| LVPW;s (mm) | 0.82±0.06 | 0.97±0.06 | 1.26±0.10 | 1.52±0.07 | 1.37±0.07 | 0.87±0.05 | 1.09±0.12 | 1.32±0.22 | 1.37±0.12 | 1.25±0.06 |
| Doppler (mm/s) |  | -3665±206 |  |  |  |  | -3671±209 |  |  |  |
| HR (bpm) | 511.23±16.33 | 513.64±11.34 | 512.55±28.10 | 506.11±26.13 | 545.55±29.02 | 470.08±12.68 | 493.20±21.93 | 492.34±24.46 | 476.12±30.90 | 498.91±19.11 |
| Body Weight (g) | 28.30±0.76 | 26.70±0.41 | 28.57±0.70 | 29.87±0.73 | 31.00±0.53 | 28.70±1.49 | 27.96±1.74 | 28.44±1.19 | 30.00±1.52 | 29.60±1.03 |
| LV Mass (mg) | 110.68±6.06 | 191.11±17.75 | 198.85±27.89 | 252.14±24.46 | 257.60±24.23 | 90.33±4.89 | 156.06±12.16 | 201.37±11.15 | 209.86±11.13 | 245.80±19.90 |
| FAC (%) | 37.78±1.55 | 27.93±3.16 | 24.25±2.52 | 23.85±1.96 | 23.38±1.37 | 46.50±1.70 | 33.50±2.68 | 31.45±4.94 | 27.06±2.83 | 22.50±3.65 |

**Supplemental Table S6.** *Atcayos* KO TAC echocardiography

| Time Point | *Atcayos*<sup>+/+</sup> |  |  |  |  | *Atcayos*<sup>-/-</sup> |  |  |  |  |
| --- | --- | --- | --- | --- | --- | --- | --- | --- | --- | --- |
|  | Baseline | Week 1 | Week 4 | Week 6 | Week 8 | Baseline | Week 1 | Week 4 | Week 6 | Week 8 |
| ENDOarea;d (mm<sup>2</sup>) | 12.95±0.65 | 15.66±1.10 | 14.89±0.89 | 15.09±1.57 | 16.15±1.51 | 14.95±1.70 | 15.21±0.94 | 14.44±0.97 | 14.17±1.26 | 15.74±1.20 |
| ENDOarea;s (mm<sup>2</sup>) | 8.11±0.43 | 10.15±0.94 | 10.71±0.84 | 11.02±1.52 | 12.47±1.38 | 8.90±0.86 | 9.95±0.81 | 10.22±0.87 | 10.65±1.25 | 12.27±1.30 |
| IVS;d (mm) | 0.77±0.03 | 1.05±0.06 | 1.08±0.05 | 1.03±0.04 | 1.08±0.05 | 0.62±0.04 | 0.90±0.09 | 1.01±0.04 | 0.99±0.05 | 1.14±0.07 |
| IVS;s (mm) | 1.01±0.06 | 1.42±0.07 | 1.32±0.06 | 1.30±0.04 | 1.39±0.06 | 0.83±0.04 | 1.33±0.05 | 1.25±0.07 | 1.18±0.05 | 1.31±0.05 |
| LVID;d (mm) | 4.01±0.09 | 4.32±0.15 | 4.47±0.14 | 4.47±0.19 | 4.45±0.20 | 4.25±0.09 | 4.22±0.15 | 4.33±0.16 | 4.24±0.17 | 4.52±0.16 |
| LVPW;d (mm) | 0.82±0.04 | 1.05±0.05 | 1.09±0.04 | 1.17±0.07 | 1.21±0.06 | 0.60±0.04 | 0.93±0.05 | 1.14±0.04 | 1.21±0.09 | 1.12±0.07 |
| LVPW;s (mm) | 0.99±0.06 | 1.24±0.06 | 1.36±0.06 | 1.36±0.03 | 1.37±0.05 | 0.77±0.05 | 1.14±0.07 | 1.35±0.06 | 1.35±0.08 | 1.33±0.05 |
| Doppler (mm/s) |  | -3457±174 |  |  |  |  | -3509±210 |  |  |  |
| HR (bpm) | 485.01±22.77 | 513.39±13.83 | 520.69±15.65 | 510.88±18.73 | 538.68±13.28 | 451.29±34.10 | 515.49±7.01 | 555.15±17.93 | 513.05±19.75 | 527.46±29.90 |
| Body Weight (g) | 28.37±0.77 | 27.40±0.51 | 28.50±0.74 | 29.31±0.59 | 29.44±0.47 | 27.36±0.81 | 26.41±0.63 | 27.26±0.74 | 28.31±0.82 | 29.29±0.84 |
| LV Mass (mg) | 117.88±7.48 | 195.39±16.13 | 216.98±14.07 | 221.22±19.53 | 234.12±20.87 | 91.61±3.64 | 154.58±11.35 | 202.07±11.93 | 203.23±18.00 | 234.41±22.46 |
| FAC (%) | 37.41±0.66 | 35.60±1.93 | 29.18±2.09 | 27.98±2.40 | 23.47±2.09 | 40.13±1.73 | 34.66±2.63 | 29.59±2.08 | 25.66±3.29 | 22.75±3.03 |

